# Spinal control of locomotion before and after spinal cord injury

**DOI:** 10.1101/2023.03.22.533794

**Authors:** Simon M. Danner, Courtney T. Shepard, Casey Hainline, Natalia A. Shevtsova, Ilya A. Rybak, David S.K. Magnuson

**Affiliations:** Department of Neurobiology and Anatomy, College of Medicine, Drexel University, Philadelphia, PA, USA; Interdisciplinary Program in Translational Neuroscience, Health Sciences Campus, Louisville, Kentucky, USA; Speed School of Engineering, Health Sciences Campus, Louisville, Kentucky, USA; Department of Neurological Surgery, Health Sciences Campus, Louisville, Kentucky, USA; Kentucky Spinal Cord Injury Research Center, University of Louisville School of Medicine, Health Sciences Campus, Louisville, Kentucky, USA

## Abstract

Thoracic spinal cord injury affects long propriospinal neurons that interconnect the cervical and lumbar enlargements. These neurons are crucial for coordinating forelimb and hindlimb locomotor movements in a speed-dependent manner. However, recovery from spinal cord injury is usually studied over a very limited range of speeds that may not fully expose circuitry dysfunction. To overcome this limitation, we investigated overground locomotion in rats trained to move over an extended distance with a wide range of speeds both pre-injury and after recovery from thoracic hemisection or contusion injuries. In this experimental context, intact rats expressed a speed-dependent continuum of alternating (walk and trot) and non-alternating (canter, gallop, half-bound gallop, and bound) gaits. After a lateral hemisection injury, rats recovered the ability to locomote over a wide range of speeds but lost the ability to use the highest-speed gaits (half-bound gallop and bound) and predominantly used the limb contralateral to the injury as lead during canter and gallop. A moderate contusion injury caused a greater reduction in maximal speed, loss of all non-alternating gaits, and emergence of novel alternating gaits. These changes resulted from weak fore–hind coupling together with appropriate control of left–right alternation. After hemisection, animals expressed a subset of intact gaits with appropriate interlimb coordination even on the side of the injury, where the long propriospinal connections were severed. These observations highlight how investigating locomotion over the full range of speeds can reveal otherwise hidden aspects of spinal locomotor control and post-injury recovery.

## Introduction

Locomotion is the essential behavior that allows animals to move in space (Orlovsky et al., 1999). Depending on the desired speed and environmental conditions, limbed animals use different gaits (Hildebrand, 1989). Rats and mice, for example, use walk and trot at slow speeds, and gallop or bound at high speeds (Cohen and Gans, 1975; Heglund and Taylor, 1988; Clarke and Still, 1999; Bellardita and Kiehn, 2015; Lemieux et al., 2016). Different gaits represent different interlimb coordination and are defined by the temporal pattern of limb movements, the phase relationships between limb pairs (Hildebrand, 1976, 1980, 1989), and the duty factor for each step cycle (the duration of the stance phase relative to the step cycle). Animals adjust these parameters to achieve stable locomotion across a range of speeds, hence making the expression of different gaits speed-dependent.

The spinal circuitry that controls locomotion includes rhythm generators for each limb located within the left and right halves of the lumbar and cervical spinal enlargements (Kato, 1990; Ballion et al., 2001; Juvin et al., 2012). These rhythm generators operate under control of supraspinal inputs and sensory feedback and define alternating flexor and extensor activities of each limb (McCrea and Rybak, 2007; Rybak et al., 2015; Kiehn, 2016). The rhythm generators are coupled via a series of commissural and long propriospinal neurons, which control the relative phase differences between them, thus defining interlimb coordination and gait (Talpalar et al., 2013; Kiehn, 2016; Ruder et al., 2016; Danner et al., 2017; Frigon, 2017). Locomotor movement and the adjustments made to interlimb coordination during speed-dependent gait transitions directly reflect the output of this circuitry. Therefore, the analysis of locomotor gaits is important for understanding the organization and operation of spinal circuitry and its descending control, both under normal conditions and following motor disorders and injuries (e.g., Shepard et al. 2022).

Spinal cord injury disrupts communication across the injury site. Thoracic injuries, as employed here, affect the long descending and ascending propriospinal interactions between limb-specific spinal circuits in the two enlargements as well as the primary ascending sensory pathways from and the primary supraspinal descending pathways to the lumbar enlargement. Studies in rodents showed that after incomplete injuries, (spontaneous) reorganization of spinal circuitry can lead to substantial recovery of locomotor function (Takeoka et al., 2014; Filli and Schwab, 2015). Thus, the analysis of locomotor behaviors before and after injury can be used to infer functional characteristics of the neural circuitry involved as well as injury-induced plasticity of this circuitry (Shepard et al., 2021, 2022).

It is important to note that the observable changes in locomotion after injury, such as speed-dependent gait expression, can be influenced by the experimental conditions. Hence, the methods used to collect locomotor data may influence the interpretation of the operation of the spinal circuitry and its involvement in post-injury recovery. For example, many studies focused on locomotion after spinal cord injury utilize treadmill-based assessments, or a Catwalk-like apparatus that naturally limits gait expression to the treadmill speed chosen or the trained behavior over short walking distances (Krizsan-Agbas et al., 2014; Fiker et al., 2020). While these approaches have the advantage of relatively long periods of stable stepping, they often do not allow researchers to obtain and analyze the full repertoire of expressed gaits or may alter the gait expression profile from what the animal would prefer if given ample distance in an overground setting. For example, in our recent experiments (Pocratsky et al., 2020), the synaptic silencing of propriospinal neurons disrupted right–left coordination at walk-trot speeds. However, we found that this disruption was context-specific: it was robustly expressed when the animal was walking on a surface with good grip (high friction) but was not observed during locomotion on the treadmill or when walking on a slick surface (low coefficient of friction).

Here, we study gait characteristics of overground locomotion with the full range of speeds in intact rats and following their recovery from either a mild-moderate contusion or a lateral hemisection injury at the T10 level of the spinal cord. We show that intact rats use a wide range of gaits that vary with speed. Following recovery from either contusion or hemisection injury, the animals recovered the ability to stably locomote over wide range of speeds, but no longer expressed the highest-speed gaits, and the maximal locomotor speed was reduced. Interestingly, animals that recovered from a mild contusion injury demonstrated the emergence of novel gaits that appear to result from maintained left–right coupling of the forelimb and hindlimb pairs together with weakened hindlimb– forelimb coupling. Analysis of the variability of limb coupling gives further insight into interlimb coordination in the experimental context where animals are trained to locomote at high speeds.

## Results

### Rats use different gaits depending on locomotor speed

To study changes in speed-dependent gait expression caused by spinal cord injury, we first determined the characteristics of overground locomotion in intact adult rats over the whole range of voluntarily generated speeds. Using food treats as incentives, rats were group-trained, two or three at a time, to traverse the entire length of a 3-m long Plexiglas tank (Figure 1A) and to complete the distance in one bout. Training was performed 3–4 days a week over a period of two weeks. Data collection was done with one animal at a time in the tank and food treats were not used. Footfall patterns of all four limbs were recorded and analyzed. A representative bout is shown in Figure 1B: the step frequency increased with the locomotor speed due to the shortening of the stance duration. The locomotor gait changed from a trot to a transverse gallop and then to a half-bound gallop, and once the half-bound gallop was expressed the instantaneous speed reached a plateau.

**Figure 1.**
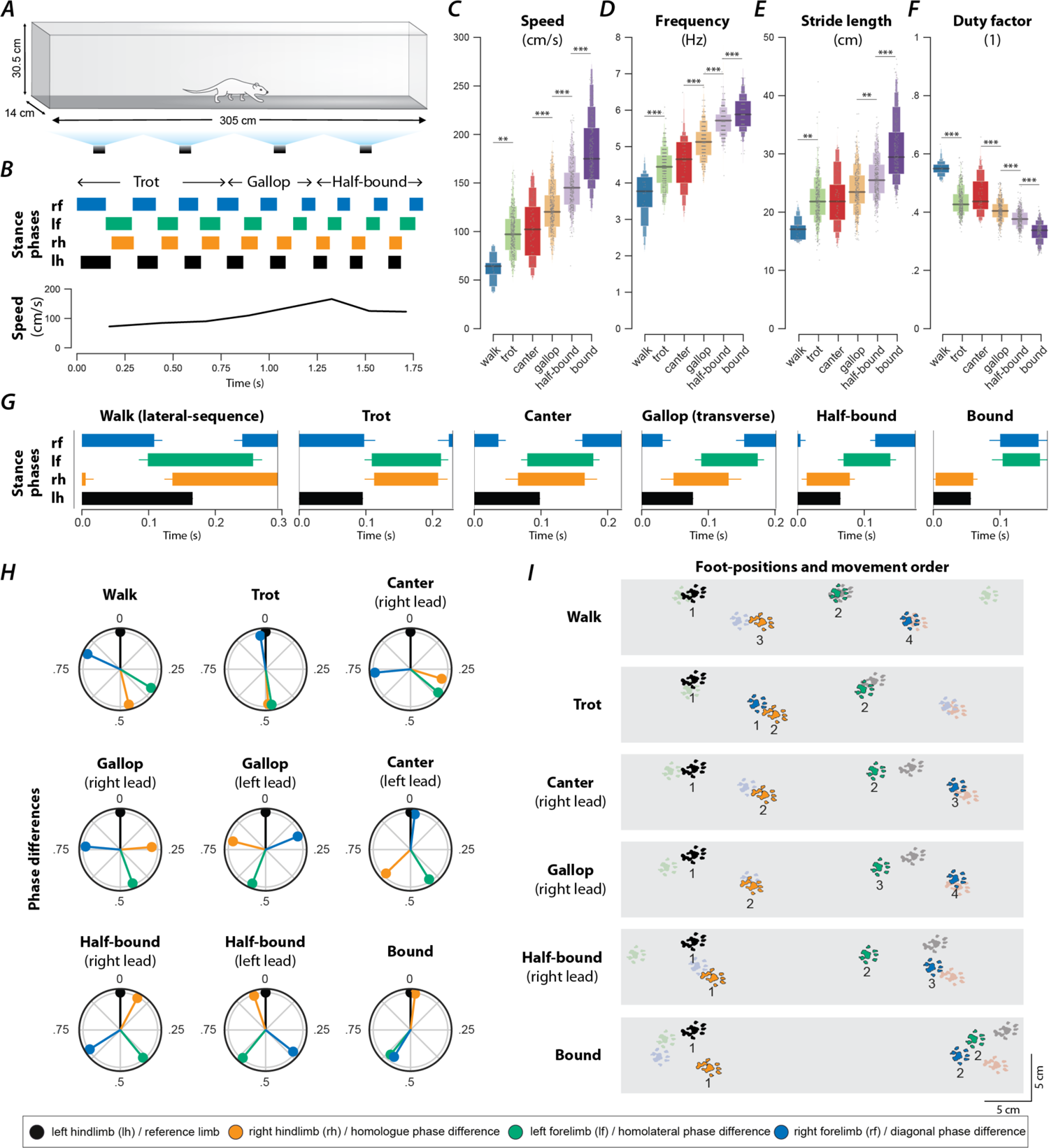
Speed-dependent gait expression in intact rats. (A) Schematic illustration of the runway chamber. (B) Stance phases and instantaneous locomotor speed of a representative bout. (C–G) Letter-value plots (Hofmann et al., 2017) of locomotor speed (C; F5,1351.4=149.166, p<0.0001), frequency (D; F5,1350.4=193.609, p<0.0001), stride length (E; F5,1353.7=68.407, p<0.0001), and duty factor (F; F5,1349.3=178.228, p<0.0001) for each gait (walk, trot, canter, gallop, half-bound and bound). The horizonal lines of the letter-value plots indicate the median, the borders of the largest boxes signify the 25^th^ and 75^th^ percentile, and the subsequently smaller boxes split the remaining data in half. For each gait parameter (C–F) a linear mixed model with gait as the fixed effect and a per-rat random intercept was calculated. Asterisks denote significant post-hoc tests between two subsequent gaits. (G) Average stance phases for each gait. (H) Circular plots of average normalized phase differences for each gait. (I) Average foot position and movement order (denoted by numbers) for each gait. **, p < 0.01; ***, p < 0.001. Detailed statistical results are reported in the Supplemental Material.

Across all animals and bouts, we identified six distinct gaits (Figure 1G–I): walk, trot, canter, transverse gallop, half-bound gallop, and bound. With the exception of canter, the mean locomotor speed (Figure 1C), step frequency (Figure 1D), and stride length (Figure 1E) increased, and duty factor (Figure 1F) decreased between gaits in the aforementioned sequence. Canter appeared to be an intermediate gait between trot and gallop where the limb pairs were not alternating and the duty-cycle dropped below 0.5, but the instantaneous speed was not different from trot. Gait transitions followed the same sequence, so that transitions between neighboring gaits in this sequence were generally most prevalent (Figure 2H1, I1). Canter was the main exception: animals were three times as likely to transition from trot directly to gallop than to canter (Figure 2H1).

**Figure 2.**
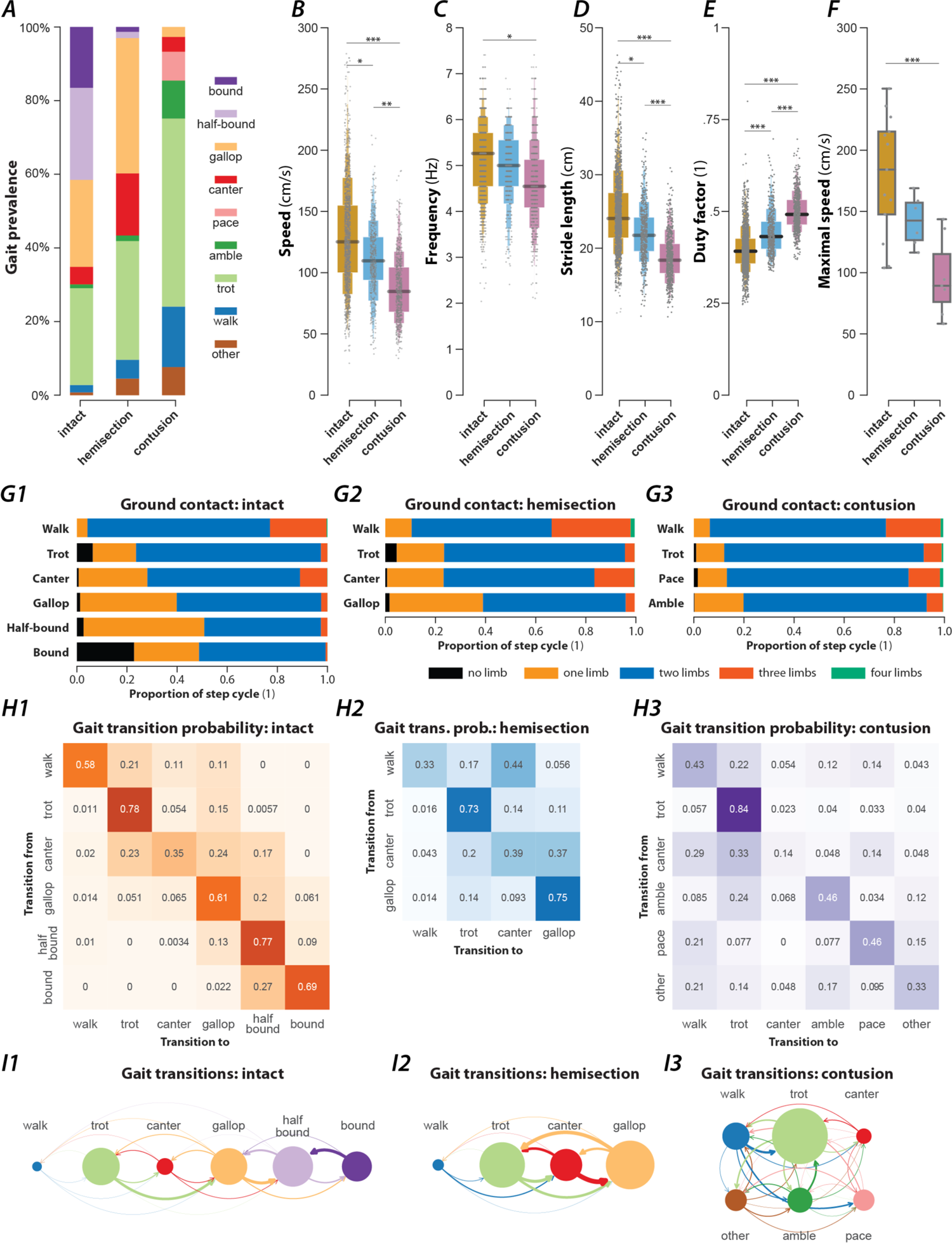
Prevalence of gaits and gait parameters in intact animals and after contusion or hemisection injury. (A) Prevalence of each gait across intact animals and animals after hemisection and contusion injury. (B–E) Boxen plots of locomotor speed [B; F2,2271=13.541, p<0.0001; intact: 127±6.83 cm/s (mean±standard error); hemisection: 110±2.62 cm/s; contusion: 85±7.17 cm/s], frequency (C; F2,2271=3.457, p=0.0317; intact: 5.08±0.14 Hz; hemisection: 4.95±0.08 Hz; contusion: 4.49±0.25 Hz), stride length (D; F2,2271=28.707, p<0.0001; intact: 24.3±0.76 cm; hemisection: 22,1±0.36 cm, contusion: 18.5±0.54 cm) and duty factor (E; F2,2271=63.225, p<0.0001; intact: 0.399±0.011; hemisection: 0.445±0.010; contusion: 0.498±0.010) for each injury type (intact, hemisection and contusion). (F) Boxplot of the maximal speeds measured as the 95^th^-percentile of the instantaneous speeds (F2,21=13.399, p=0.0002; intact: 188±8.85 cm/s, hemisection: 156±12.87 cm/s, contusion: 119±11.93 cm/s). (G) Number of limbs on the ground as a proportion of the step cycle for each gait (G1: intact, G2: hemisection, G3: contusion). (H) Matrices of gait transition probabilities (H1: intact, H2: hemisection, H3: contusion). (I) Gait transition graphs (I1: intact, I2: hemisection, I3: contusion), where nodes represent gaits (size is proportional to their prevalence) and edges represent gait transitions (line widths are proportional to their frequency of occurrence). For each gait parameter (B–F) a mixed model with injury type as the fixed effect, a per-rat random intercept and slope of injury type was calculated. For speed and frequency, linear models were calculated; for stride length, a log link was used; for duty factor, a Gamma error distribution and a logit link function was used. Asterisks denote significant post-hoc tests between two subsequent gaits. *, p < 0.05, **, p < 0.01; ***, p < 0.001. Detailed statistical results are reported in the Supplemental Material.

### Walk

Walk is a four-beat gait with longer stance than swing durations (duty factor is greater than 0.5). Walks can be subdivided into six groups based on the sequence of steps. In our study, the rats mainly used a lateral-sequence walk pattern (75.9% of all walking steps, n=22 steps, 8 of 17 rats), in which a hindlimb step was followed by that of the ipsilateral forelimb, the contralateral hindlimb, and then the contralateral forelimb (Figure 1G). The hindlimb paw placements were close to those of the ipsilateral forelimb in the previous step cycle (Figure 1I). Compared to all other gaits (Figure 1C–F), walk was expressed at the lowest locomotor speeds (77.1±6.56 cm/s) and step frequencies (3.98±0.12 Hz), had the shortest stride length (18.8±0.97 cm), and highest duty factor (0.52±0.01). At any time, either two or three limbs were on the ground simultaneously (Figure 2G1).

It is not surprising that very few steps were categorized as a walk (2%, n=29; Figure 2A) because the animals were trained to locomote quickly from one end of the runway to the other. The average duty factor was close to 0.5 with homolateral and diagonal phase differences between those of an ideal lateral-sequence walk and those of trot (see Table 1; if not otherwise noted, the phase differences are normalized to the range of [0,1), representing the delay between midstance times of a pair of limbs divided by the step cycle duration). This suggests that walk was mainly used as a transitory gait at the beginning of a bout.

**Table 1.**
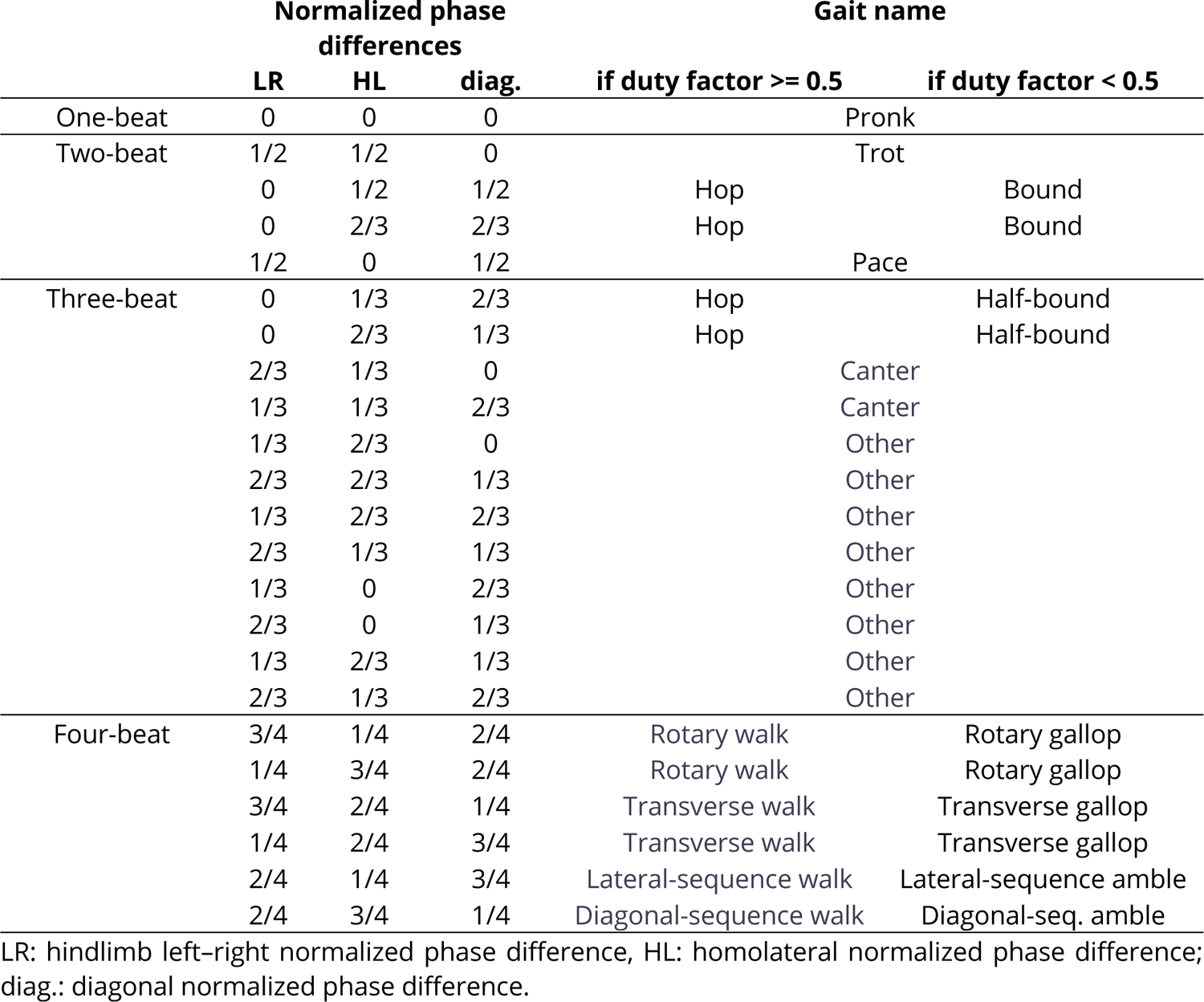
Idealized gaits

### Trot

Trot is characterized by synchronous (in-phase) movements of the diagonal limb pairs (normalized diagonal phase difference close to 0 or 1) and alternation of the movements of all other pairs of limbs (phase differences ∼0.5). Trot was expressed at an intermediate speed (100.5±3.61 cm/s; Figure 1C), frequency (4.46±0.07 Hz; Figure 1D), and stride length (22.2±0.46 cm; Figure 1E), and with a duty factor well below 0.5 (0.43±0.007; Figure 1F). Since the diagonal limbs moved in synchrony, the duty factor below 0.5 resulted in a brief suspension phase (no limbs on the ground) between the lift-off of one pair and the touchdown of the other pair of diagonal limbs (Figure 1G, Figure 2G1). Thus, the observed gait was a “running trot” (Górska et al., 1999). The relative footfall positions were comparable to those used in the lateral-sequence walk: each hindlimb closely approximated the position of its ipsilateral forelimb. Trot was the most prevalent (26.3%, n=381, 16/17 rats; Figure 2A) and stable gait (Figure 2H1), which is in accordance with previous reports in rats and mice (Cohen and Gans, 1975; Lemieux et al., 2016).

#### Canter

Canter is a three-beat gait in which one pair of diagonal limbs moves in synchrony. The footfall of the synchronously moving diagonal limbs is preceded by that of the other hindlimb and followed by that of the other forelimb. The leading forelimb is the one that is not part of the synchronous diagonal pair. Note that the leading limb is defined as the second limb of a couplet (fore or hindlimb) to touch down, which is also the limb that advances further forward within one step-cycle (Hildebrand 1977). Canter (Figure 1G–I) was the second least prevalent gait (5.8%, n=69, 15/17 rats; only walk occurred less often; Figure 2A) and the least stable one (only 33% of canter steps were followed by another step of canter; Figure 2H1). It is interesting to note that almost no steps (0.2%, n=3) with the reversed sequence (a forelimb followed by the diagonal pair followed by the hindlimb) were observed. The canter observed in our study did not exhibit a suspension phase (Figure 2G1)—a common feature of canter observed in other species such as dogs and horses (Harris, 1993; Bertram, 2016). The average speed (106.3±4.84; Figure 1C), frequency (4.65±0.09 Hz; Figure 1D), stride length (22.4±0.68 cm; Figure 1E), and duty factor (0.44±0.008; Figure 1F) of the canter were similar to trot. As the gait shares characteristics of trot (diagonal synchronization) and gallop (left–right asymmetry), it likely occurred at the transition between (Figure 2H1, I1) them in preparation for increased speed exhibited during gallop.

#### Gallop

Gallop represents a class of left–right non-alternating, four-beat gaits, in which the forelimbs and hindlimbs move in pairs but with some delay between the lead and trailing limb of each pair. Gallops can be subdivided into transverse gallops and rotary gallops based on the footfall sequence. If the forelimb pair has the same lead limb (left or right) as the hindlimb pair, the sequence is referred to as transverse; if the forelimb and hindlimb pairs have the opposite lead limb, the sequence is referred to as rotary. In our study, the transverse gallop was dominating (23.5%, n=340, 17/17 rats; Figure 1G– I) while the rotary gallop was observed in only a few episodes (0.14%, n=2, 2/17 rats). The transverse gallop was expressed on average at higher speed (122.4±3.62 cm; Figure 1C), frequency (5.11±0.074 Hz; Figure 1D), and stride length (23.8±0.457 cm; Figure 1E), and at a lower duty factor (0.40±0.007; Figure 1F) than trot. The footfall pattern of the four limbs was evenly spaced throughout the gait cycle and either one or two limbs were on the ground at any time; there was no suspension phase (Figure 2G1).

#### Half-bound gallop

Half-bound gallop is a special case of gallop; it is a non-alternating three-beat gait in which the hindlimbs, but not the forelimbs, move in synchrony. Thus, it is distinct from the other gaits because the two limb pairs have different phase differences. The half-bound gallop (Figure 1G–I) was similar in prevalence (25%, n=362, 16/17 rats; Figure 2A) to trot and transverse gallop, and one of the most stable gaits (Figure 2H1). It is a high-speed gait and was expressed at higher speeds (141±3.62 cm/s; Figure 1C), frequencies (5.57±0.07 Hz; Figure 1D), and stride lengths (25.1±0.46 cm; Figure 1E) than the transverse gallop. The average duty factor was 0.38±0.007 (Figure 1F) and, similarly to the transverse gallop, the half-bound rarely exhibited a suspension phase; one or two limbs were on the ground at all times (Figure 2G1).

#### Bound

Bound is a non-alternating gait characterized by in-phase (synchronous) movements of both the hindlimb and the forelimb pairs. Hindlimb and forelimb stance phases were separated by a suspension phase (Figure 1G; Figure 2G1), however the suspension phase following hindlimb stance was longer than the one following forelimb stance. The hindlimb pair touched down in front of the forelimb positions of the previous step cycle (Figure 1I). Bound occurred in 16.5% of the steps (n=239, 12/17 rats) and hence was the fourth most prevalent gait (Figure 2A). It was expressed at the highest locomotor speeds (173.9±3.94 cm/s; Figure 1C), frequencies (5.73±0.08 Hz; Figure 1D), longest stride lengths (30.0±0.52 cm; Figure 1E), and smallest duty factors (0.34±0.007; Figure 1F).

Very few steps (1.8%, n=26, 4/17 rats) were categorized as belonging to any other gait (see Table 1), such as pronk, pace or amble, hop, diagonal-sequence walk, rotary-sequence gallop, and other three-beat gaits (besides canter and half-bound gallop).

### Rats recovered from spinal cord injury exhibited a reduced locomotor speed

To study gait characteristics after spinal cord injury, a mid-line T9 contusion or a lateral T9 hemisection was performed in subsets of the intact animals (hemisection: 6 rats; contusion: 7 rats). These injuries result in grossly similar chronic locomotor deficits as assessed by the BBB Open Field Locomotor Scale, with mean scores of 15.7±1.9 (contused) and 18.3±0.8 (hemisected) on the scale of 0–21 at day 42 post-injury. Thus, the injuries allowed for recovery sufficient for gait analysis with consistent plantar paw placement (no dorsal steps), and the appearance of forelimb–hindlimb coordination despite substantial loss of ascending-descending axons (Magnuson et al., 2005; Ballermann and Fouad, 2006) Importantly, hemisected animals have significant right-left differences in hindlimb function acutely post-injury, and the spontaneous recovery of hindlimb function ipsilateral to the injury is known to be accompanied by significant sprouting of spared descending motor tracts below the level of injury (Fouad et al., 2000; Ballermann and Fouad, 2006). In contrast, animals with mid-line contusions show largely symmetrical recovery and the role played by sprouting below the injury is less clear (Magnuson et al., 2005; Fink and Cafferty, 2016). Injured animals were re-introduced to the plexiglas tank and re-trained, again as a small group, over a period of four days, during week five post-injury, using food treats as incentive to fully traverse the walkway in one bout. Characteristics of their recovered locomotor function were then assessed using the same methods as those used before the injury. Exemplary bouts of a rat with a lateral hemisection injury or a contusion injury can be seen in Figure 5A and Figure 7A.

The rats in each injury group exhibited reduced mean locomotor speeds compared to pre-injury (Figure 2B), with post-contusion animals having a lower mean speed than those with a hemisection. Interestingly, the maximum speed (measured as the 95^th^-percentile of the instantaneous speed) was significantly reduced after contusion but not after hemisection injury (Figure 2F).

Like in other limbed animals, locomotor and gait parameters in rats in our experiments depended on and changed with locomotor speed (Figure 3). Specifically, the stride length increased linearly with speed (Figure 3A), while the step-cycle duration (or period) showed an inversely proportional decrease (Figure 3B)—thus the step frequency increased with speed—and the duty factor decreased linearly with speed (Figure 3C). This speed-dependent change of the duty factor was caused by an asymmetric change of the stance and swing durations with speed: the stance duration was inversely related to speed (Figure 3D), while the swing duration only decreased linearly with speed (Figure 3E); the change with speed was larger for the stance duration than for the swing duration.

**Figure 3.**
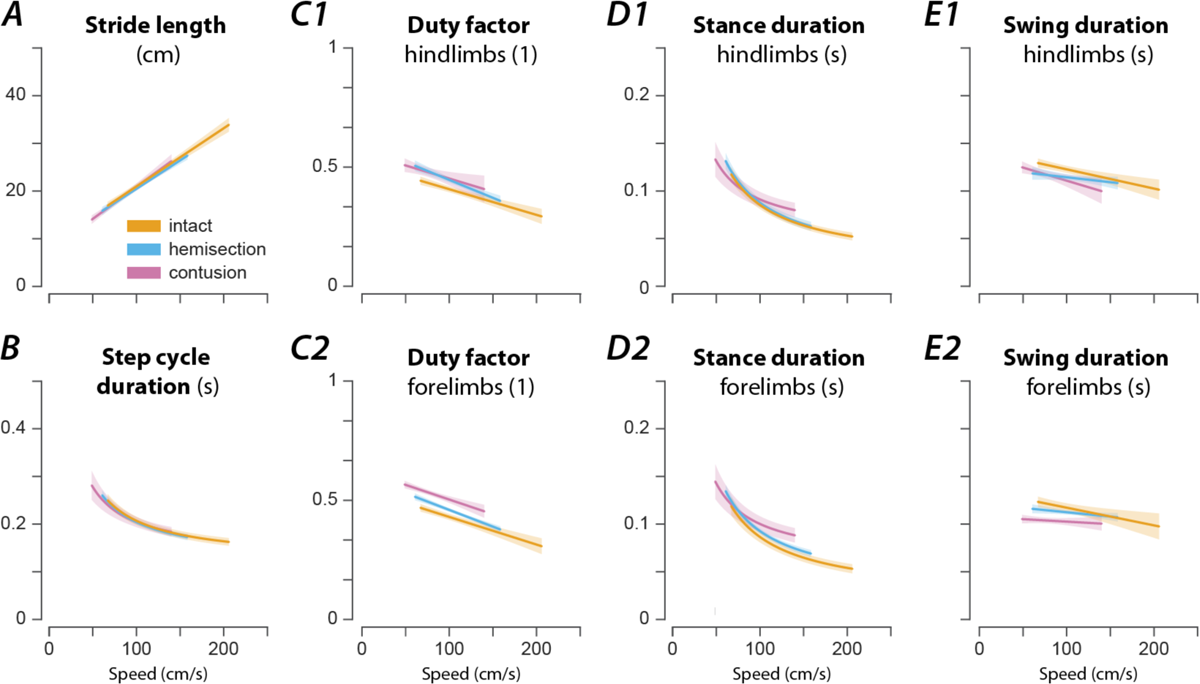
Speed-dependent modulation of gait characteristics in intact rats and after hemisection and contusion injury. Regression plots of stride length (A), step cycle duration (B), hindlimb (C1) and forelimb duty factor (C2), hindlimb (D1) and forelimb stance duration (D2), and hindlimb (E1) and forelimb swing duration (E2) versus speed. Solid lines represent the regression result and the shaded areas the 95%-confidence intervals. For each parameter, regressions were calculated for intact rats (orange) and after hemisection (blue) and contusion injury (purple). Linear regressions were calculated for the stride length (A), duty factors (C) and swing duration (E); non-linear regressions with the function β_0_ + β_1_ * ν^-1^, where β_0_ and β_1_ are regression parameters and ν the instantaneous speed, were calculated for the step cycle duration (B) and stance durations (D). All regressions were calculated on a per-animal basis and then averaged. Regression parameters and results of post-hoc comparisons are presented in the Supplementary Material.

After either spinal cord injury, the speed-dependent gait characteristics remained qualitatively similar to pre-injury with only minor differences (Figure 3). The changes in stride length (Figure 3A) and step cycle duration (Figure 3B) with speed were not significantly different from the pre-injury values. The forelimb and hindlimb duty factors (Figure 3C) after contusion injury increased with the speed across the expressed speed range with significantly increased offsets; after hemisection injury the duty factors had a significantly larger negative slope than before the injury, resulting in higher duty factor values than before the injury at low speeds but not at high speeds. Furthermore, contused animals showed significant changes in the modulation of the forelimb and hindlimb stance durations (Figure 3D) and a significantly decreased forelimb swing duration (Figure 3E). Hemisected animals showed a significant reduction of the modulation of the swing duration (Figure 3E) and demonstrated a significant increase in forelimb stance duration compared to the intact case (Figure 3D2). Finally, after hemisection, there was a left–right asymmetry of hindlimb, but not forelimb duty factors (Figure 4): the duty factor of the left hindlimb (contralesional) was longer than that of the right (ipsilesional) hindlimb.

**Figure 4.**
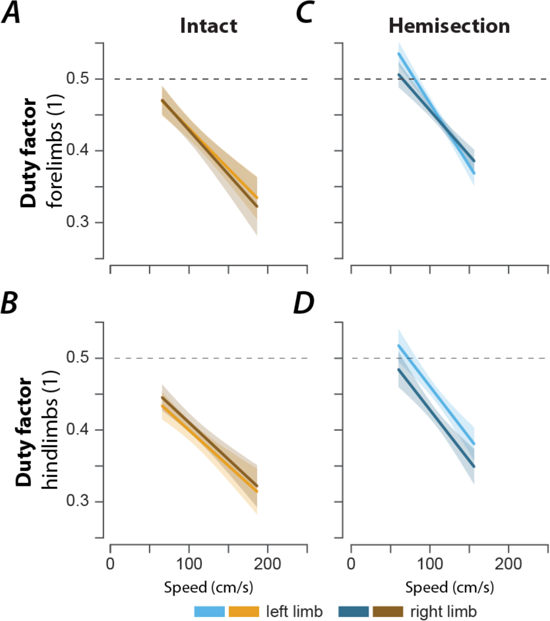
Left–right asymmetry of duty factors after right lateral hemisection. Regression plots of left and right forelimb (A, C) and hindlimb duty factors (B, D) versus speed for intact (A, B) and hemisected animals (C, D). Shaded areas represent 95% confidence intervals. Hemisection was applied to the right side of the spinal cord.

### Rat recovered from hemisection could not move with high-speed synchronous gaits and showed a lead limb bias during asymmetric gaits

In the exemplary locomotor bout shown in Figure 5A, the hemisected rat started with a walk, then, while accelerating, transitioned to trot, followed by an intermediary step of canter and two steps of gallop. At the end of the bout, the rat decelerated and transitioned back to a trot. Throughout the bout, it used neither half-bound gallop nor bound—the highest speed gaits observed before the injury.

**Figure 5.**
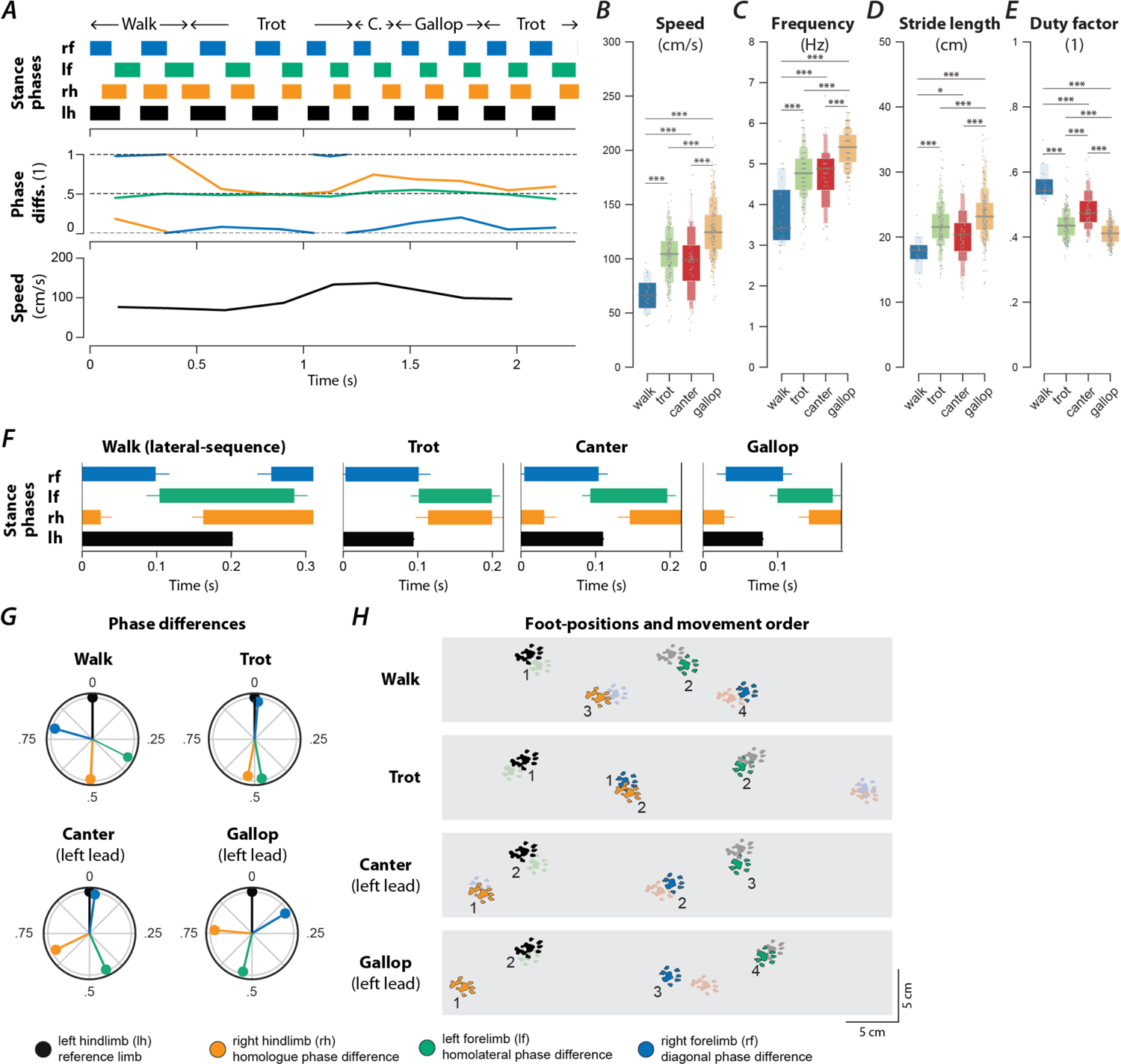
Speed-dependent gait expression after recovery from a right lateral hemisection injury. (A) Stance phases, normalized phase differences, and instantaneous locomotor speed of a representative bout. (B–E) Boxenplots of locomotor speed (B; F3,402=83.522, p<0.0001), frequency (C; F3,103.42=70.811, p<0.0001), stride length (D; F3,406.17=22.221, p=<0.0001), and duty factor (E; F3,44.88=94.177, p<0.0001) for each gait (walk, trot, canter, and gallop). The horizonal lines of the letter-value plots indicate the median, the borders of the largest boxes signify the 25^th^ and 75^th^ percentile, and the subsequently smaller boxes split the remaining data in half. For each gait parameter (B–E) a linear mixed model with gait as the fixed effect and a per-rat random intercept was calculated. Asterisks denote significant pairwise post-hoc tests. (F) Average stance phases for each gait. (G) Circular plots of average normalized phase differences for each gait. (H) Average foot position and movement order (denoted by numbers) for each gait. C., canter; *, p < 0.05; **, p < 0.01; ***, p < 0.001. Detailed statistical results are reported in the Supplemental Material.

Across all animals and locomotor bouts (Figure 2A), half-bound gallop [1.7%, n=8, odds ratio (OR) compared to before injury=0.07, 5/6 rats] and bound (1.3%, n=6, OR=0.08, 2/6 rats) were essentially absent. The loss of these high-speed gaits was compensated for by an increased prevalence of gallop (36.7%, n=172, OR=1.56, p<0.0001, 6/6 rats, Fisher’s exact test), canter (16.9%, n=79, OR=3.54, p<0.0001, 6/6 rats, Fisher’s exact test), walk (5.1%, n=24, OR=2.56, p=0.0009, 5/6 rats, Fisher’s exact test), and trot (33.3%, n=151, OR=1.23, p=0.0149, 6/6 rats, Fisher’s exact test). As in the intact case, these gaits were expressed at different locomotor speed (Figure 5B), frequency (Figure 5C), stride length (Figure 5D), and duty factor (Figure 5E). Walk was expressed at the lowest average speed, frequency and stride length, trot and canter at intermediate values, and gallop at the highest values; the average duty factor sequentially decreased from walk to canter and then to trot and gallop. Note that the average values of speed, frequency and stride length did not differ between trot and canter, similar to the intact condition. Trot and gallop were the most stable gaits (Figure 2H2) and, similar to the intact case, canter was mostly expressed when the animals transitioned between trot and gallop (Figure 2H2, I2). Finally, the hemisected rats did not express gaits that were not observed before the injury.

Interestingly, the leading forelimb (the second forelimb to touch down, which is also the forelimb that advances forward to a greater extent; Hildebrand 1977) during asymmetric gaits (canter and gallop) was almost exclusively the left one across all hemisected animals (Figure 6A,B). This was a significant change compared to locomotion before injury, where the animals used the left forelimb as a leading limb in roughly one third of the step cycles. As the lateral hemisection injury was performed on the right side of the spinal cord, the lead limb preference shifted to the side contralateral to the injury. Thus, the limbs ipsilateral to the injury touched down first and the limbs contralateral to the injury touched down second (lead limb). This was the case for both the forelimbs and hindlimbs because transverse gallops (the only gallop expressed) and canter used the same lead for forelimbs and hindlimbs. Thus, the hemisection caused a clear left–right bias of the limb coordination during asymmetric gaits that had the same directionality in the forelimbs and hindlimbs.

**Figure 6.**
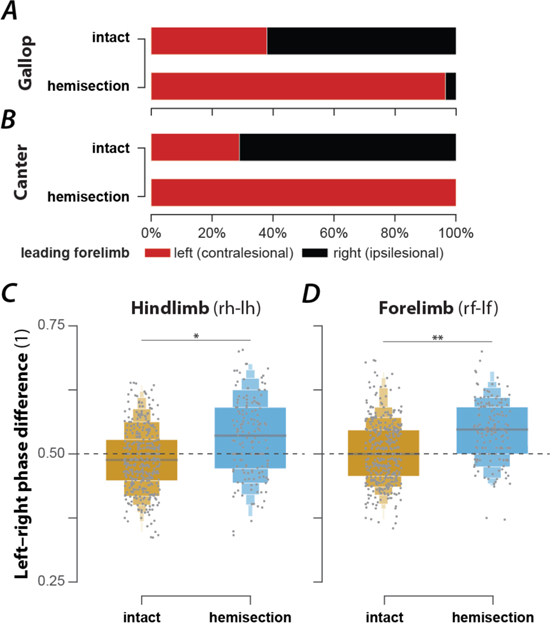
Changes of lead-limb preference (A, B) and left–right coordination during trot (C, D) following recovery after right lateral hemisection injury. (A–B) Prevalences of lead-limbs (left or right forelimb that touches down second) pre-injury (intact) and after hemisection for gallop (A; intact: 37.4%, n=127; hemisection: 96.5%, n=166; p<0.0001; Fischer’s exact test) and canter (B; intact: 29.0%, n=20; hemisection: 100%, n=79; p<0.0001; Fischer’s exact test). (C–D) Boxenplots of hindlimb (C; intact: 0.493±0.009; hemisection: 0.549±0.017; *X*^2^_1_=5.962, p=0.0146) and forelimb normalized left–right phase differences (D; intact: 0.508±0.006; hemisection: 0.546±0.006; *X*^2^_1_ =9.953, p=0.0016). (C–D) For both hindlimb and forelimb normalized phase differences a linear mixed model with injury type as a fixed effect, a random per-animal intercept, a random slope of injury type, and a full dispersion model were calculated. l, left; r, right; f, forelimb; h, hindlimb, **, p < 0.01; ***, p < 0.001.

To test whether the hemisection injury also caused a left–right bias of interlimb coordination during alternating gaits, we compared the forelimb and hindlimb left–right phase differences for trot between hemisected rats and intact rats (Figure 6C,D). The means of both left–right phase differences were significantly increased in hemisected animals compared to the intact ones. Thus, after hemisection, movements of the homologous limbs were not perfectly alternating during trot; instead, there was a bias so that the movements of the left (contralesional) limbs followed those of the right (ipsilesional) limbs more quickly than vice versa. Thus, even for the alternating gait of trot, the limbs contralateral to the hemisection became the lead with a slight asymmetry in the alternation of each limb pair.

In summary, lateral hemisection resulted in a loss of the high-speed synchronous gaits (half-bound gallop and bound) and caused a left-right bias in the remaining gaits.

### Contusion results in a loss of high-speed gaits and the emergence of new gaits

An exemplary bout of locomotion of a rat with a contusion injury is shown in Figure 7A. The animal starts with lateral-sequence walk that then transitions to two steps of pace (characterized by in-phase movements of each homolateral pair of limbs and left-right alternation of the forelimbs and hindlimbs) followed by a step of diagonal-sequence walk before it transitioned to a trot and finally back to a lateral-sequence walk. The instantaneous speed was highest during the pace steps but otherwise showed little modulation and was generally lower than that of the intact animal (Figure 1B).

**Figure 7.**
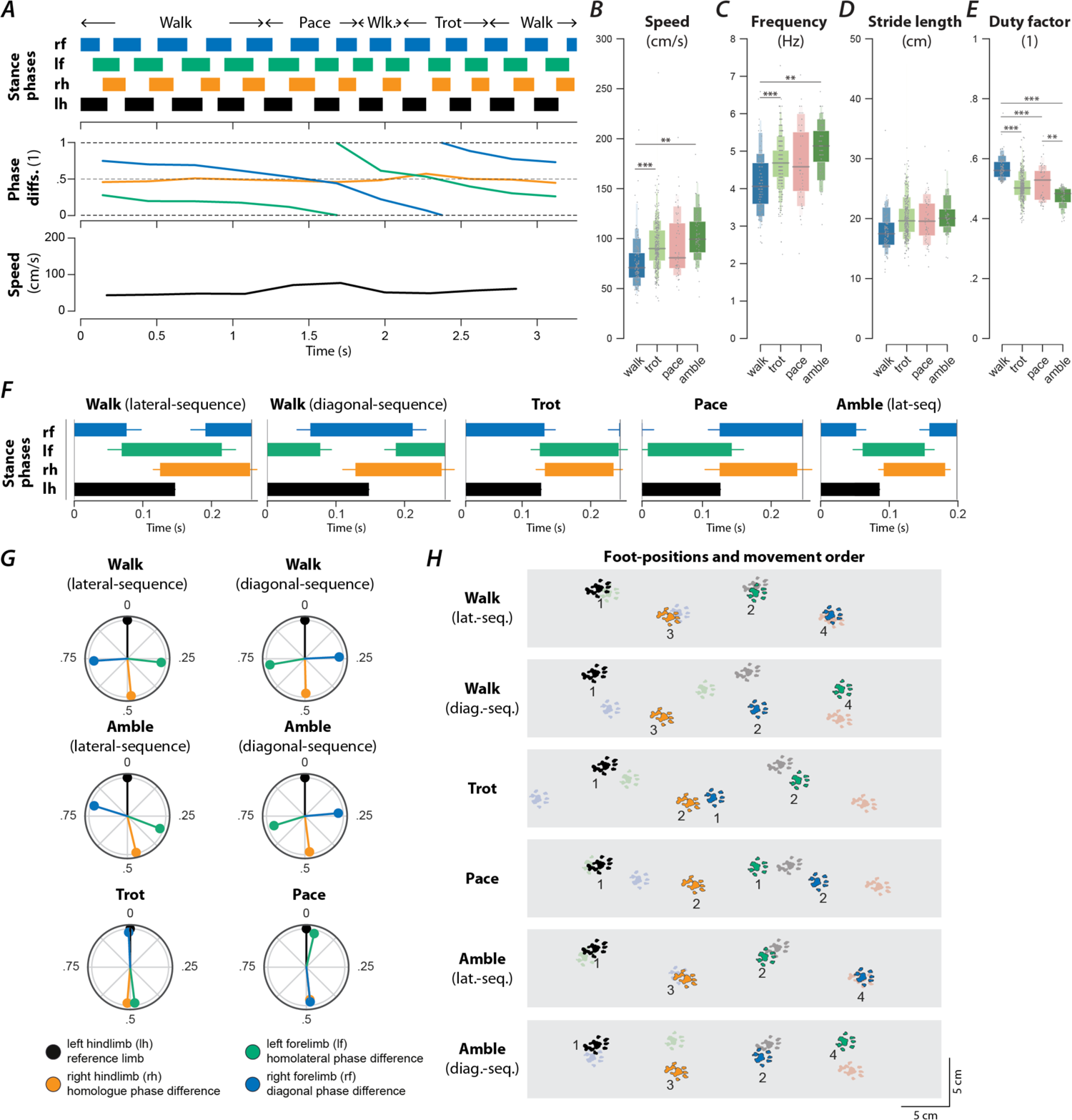
Speed-dependent gait expression after recovery from a contusion injury. (A) Stance phases, normalized phase differences, and instantaneous locomotor speed of a representative bout. (B–E) Boxenplots of locomotor speed (B; F3,531.42=9.140, p<0.0001), frequency (C; F3,530.65=10.728, p<0.0001), stride length (D; F3,492.95=2.346, p=0.0720), and duty factor (E; F3,531.57=55.583, p<0.0001) for each gait (walk, trot, canter, and gallop). The horizonal lines of the letter-value plots indicate the median, the borders of the largest boxes signify the 25^th^ and 75^th^ percentile, and the subsequently smaller boxes split the remaining data in half. For each gait parameter (B–E) a linear mixed model with gait as the fixed effect and a per-rat random intercept was calculated. Asterisks denote significant pairwise post-hoc tests. (F) Average stance phases for each gait. (G) Circular plots of average normalized phase differences for each gait. (H) Average foot position and movement order (denoted by numbers) for each gait. **, p < 0.01; ***, p < 0.001. Detailed statistical results are reported in the Supplemental Material.

Across animals (Figure 2A), the prevalence of the slow-speed alternating gaits, walk (16.4%, n=105, OR=8.19, p<0.0001, 7/7 rats, Fisher’s exact test) and trot (51.1%, n=327, OR=1.94, p<0.0001, 7/7 rats, Fisher’s exact test), increased compared to pre-injury. Indeed, trot was used in the majority of step cycles after contusion injury. None of the contused animals expressed any steps that were classified as bound or half-bound gallop, and very few steps of gallop were expressed (2.7%, n=17, OR=0.11, p<0.0001, 4/7 rats, Fisher’s exact test). Thus, the contusion resulted in a loss of high-speed non-alternating gaits.

Furthermore, the contused rats used gaits that were not (or very rarely) observed before injury: amble (10.3%, n=66, OR=9.96, p<0.0001, 7/7 rats, Fisher’s exact test), pace (7.8%, n=50, OR=οο, 6/7 rats), and diagonal-sequence walk (5.2%, n=33, OR=οο, 7/7 rats). Average stance phase duration and phase differences of the new gaits are shown in Figure 7F–G. Note that the ambling gaits (four-beat gaits comparable to walk but with a duty factor lower than 0.5) exhibited by the contused animals used both a lateral (6.0%, n=38, 6/7 rats) and a diagonal sequence (4.4%, n=28, 6/7 rats). The commonality between the novel gaits is that they exhibit left–right alternation of both the forelimb and hindlimb pairs (fore and hind left–right normalized phase difference around 0.5; see Table 1). Indeed, a large majority of the total steps (85.6%, n=548) exhibited left–right alternation—because trot and lateral-sequence walk are also left–right alternating gaits. Yet, the gaits span the whole spectrum of homolateral and diagonal phase differences.

Walk was expressed at lower average speeds and frequencies than trot and amble (Figure 7B,C), and exhibited a greater duty factor than all other gaits (Figure 7E). Yet, there were no significant differences in the average stride lengths between the gaits (Figure 7D). Furthermore, ordered gait transitions, as was the case for intact (Figure 2H1, I1) and hemisected rats (Figure 2H2, I2), appeared to be absent and gait transitions occurred between most pairs of gaits (Figure 2H3, I3).

In summary, the contusion injury caused a loss of left–right asymmetric and left–right synchronous (non-alternating) gaits, while allowing for gaits with various modes of forelimb–hindlimb coordination. These observations suggest that the modulation of stepping required for the non-alternating gaits may be dependent on the long-propriospinal neurons that interconnect the forelimb circuits with those of the hindlimbs.

### Gaits are distributed on a continuum in phase-space

Gaits are mainly defined by interlimb coordination, which is quantitatively represented by a set of phase differences between pairs of limbs. Interlimb coordination, and consequently gait characteristics, choices and transitions are controlled by spinal cord circuitry. Thus, the distribution of phase differences for each limb pair and how they change with speed provides insight into the underlying neuronal control and changes in that circuitry following spinal cord injury.

In intact rats (Figure 8A), the speed-dependent gait transitions from trot through gallop and half-bound gallop to bound can be seen as speed-dependent changes in the normalized phase differences. At low speeds (∼50–75 cm/s), left–right alternation of the hindlimbs and the forelimbs dominated. At medium speeds (∼75–150 cm/s), left–right phase differences showed progressive deviation from alternation in both directions up to synchronization (in-phase movements), while some steps still maintained left–right alternation. Finally, at the highest speeds (>150 cm/s), steps with left– right alternation were essentially absent, and most steps exhibited normalized left–right phase differences close to 0 or 1 (in-phase movements). The speed-dependent changes in the left–right phase differences in the forelimb and hindlimb were qualitatively similar, but the forelimbs rarely reached perfect synchrony. The normalized diagonal phase differences followed a similar pattern but were offset by approximately half a cycle. The homolateral phase differences remained alternating across speeds but changed from slightly below 0.5 (forelimb follows homolateral hindlimb) to just above 0.5 (hindlimb follows homolateral forelimb).

**Figure 8.**
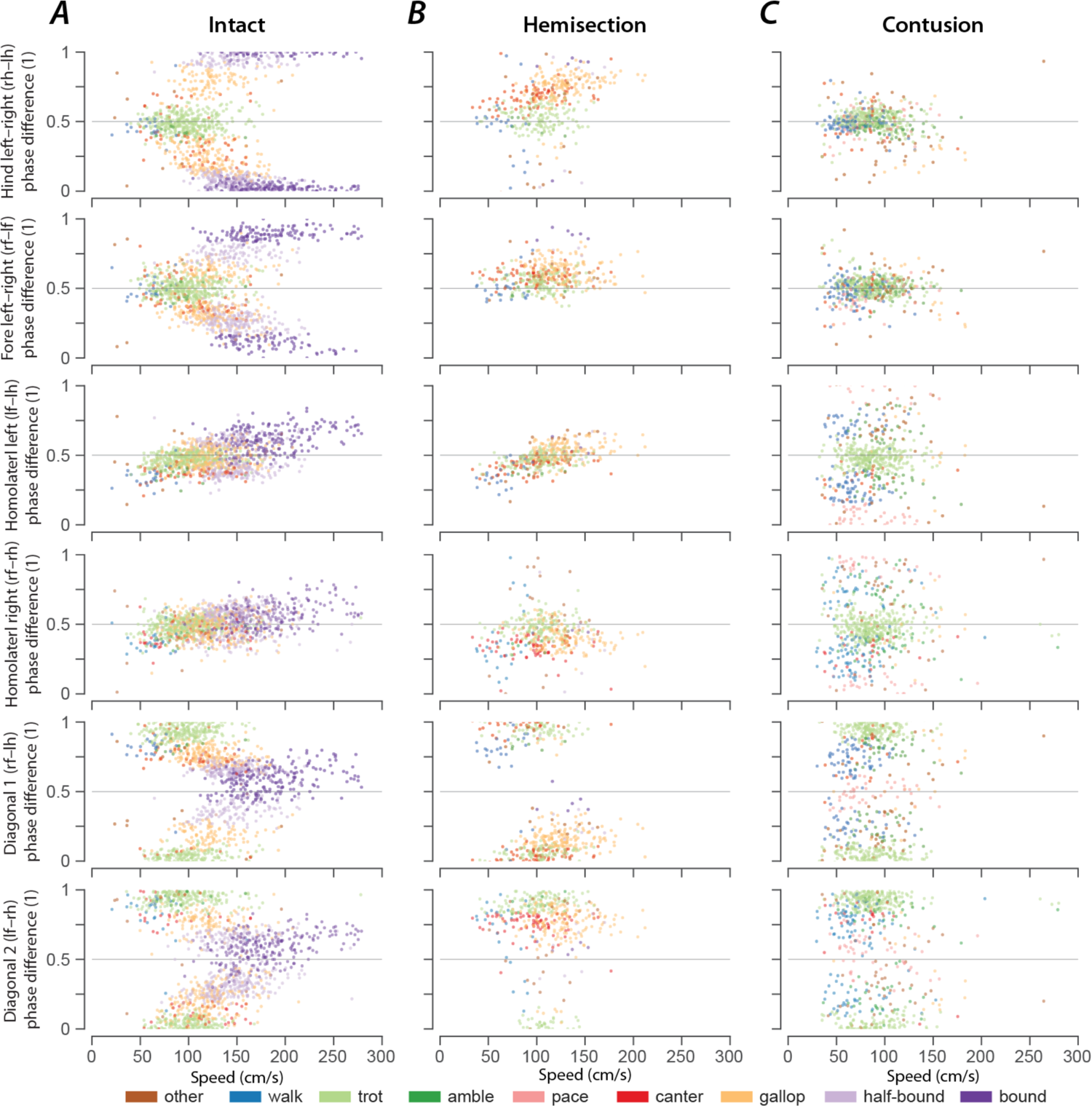
Speed-dependent distribution of normalized phase differences of intact (A), hemisected (B), and contused (C) animals. 2-dimensional scatter plots of normalized phase differences against locomotor speed. Each dot represents one step cycle. The color of the dot denotes the locomotor gait. A normalized phase difference of 0.5 signifies alternation; a normalized phase difference of 0 or 1 signifies synchronization of the two corresponding limbs. l, left; r, right; f, forelimb; h, hindlimb.

The similarity of the speed dependent changes of both left–right phase differences and the diagonal phase differences (half-cycle shift) suggests low dimensionality of the structure in phase-space and potentially in the underlying neuronal control. Indeed, the pair-wise scatterplots of normalized phase differences independent of speed (Figure 9A) reveal that most step cycles were within a small area of the space of possible phase differences. All steps of the intact rats were distributed along a line covering the range of the expressed gaits. While not completely straight, the distribution can be approximated by a line from the average trot to the average half-bound gallop with a left lead limb and its mirrored counterpart with a right lead limb (black line in Figure 9A). This model accounts for 93.72% of the variance across animals (R^2^ per rat: median=0.914, inter quartile range: 0.805–0.931). This is specifically interesting because there are several different forms of gallop that the animals could express, but all gallop steps generated fell on the line between trot and half-bound gallop—this includes gallops with either right or left limb as lead. These results suggest that the same underlying control mechanisms govern the gait changes from trot to gallop and half-bound gallop, and that these might not be discrete gaits, but are rather expressed as a continuum.

**Figure 9.**
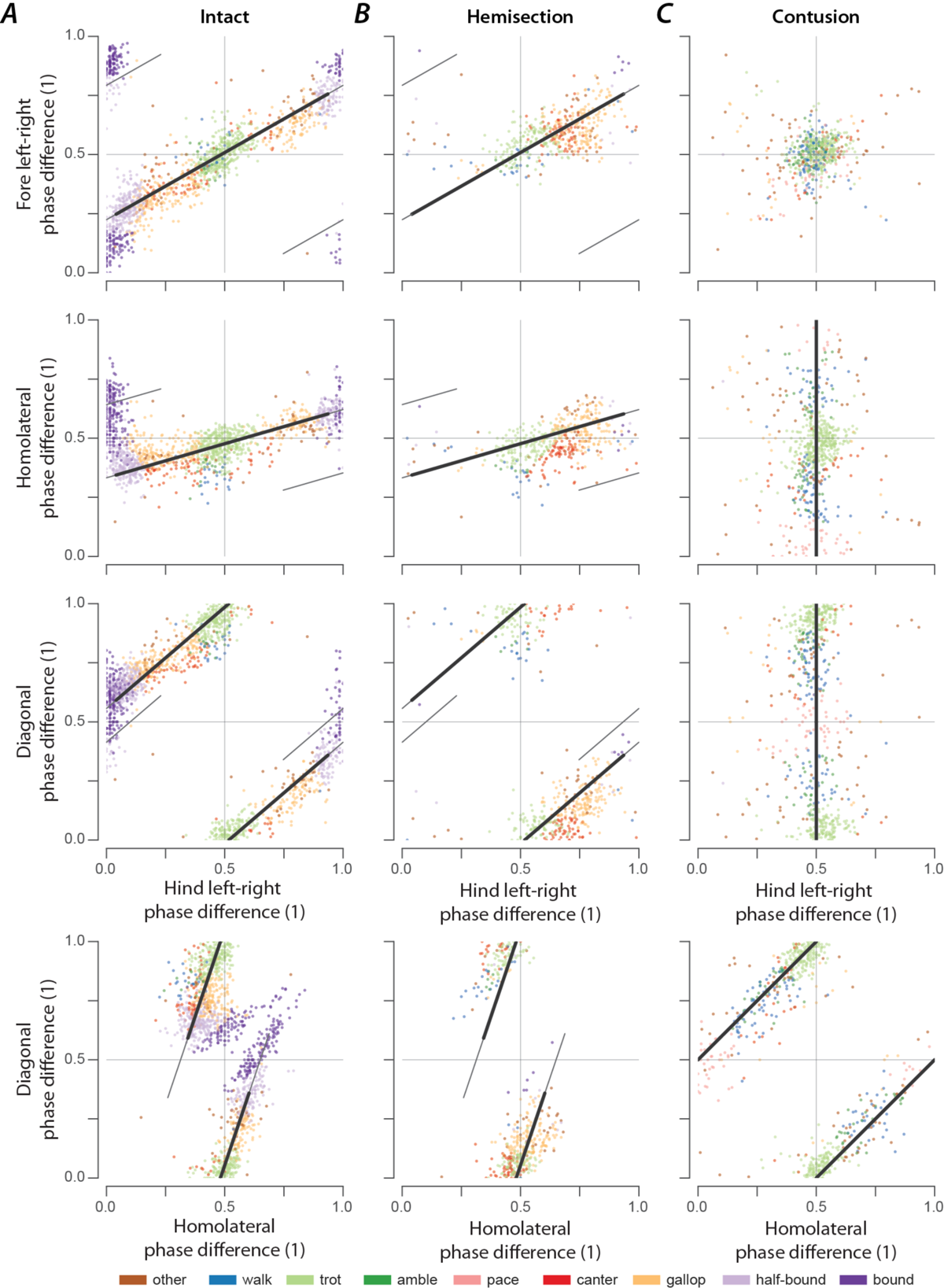
Distribution of phase differences in intact rats (A) and after hemisection (B) and contusion injury (C). (A–C) Two-dimensional scatter plots of pairs of normalized phase differences. Each dot represents a step cycle. The solid black lines span from the average trot [Ψ_A_ = (Ψ_LR_, Ψ_FH_, Ψ_DIAG_) = (0.49,0.47,0.98)] to the average half-bound gallops [Ψ_B_ = (Ψ_LR_, Ψ_FH_, Ψ_DIAG_) = (0.94,0.60,0.36)] for the intact case (A) and for the hemisection (B) and from the idealized trot [Ψ_A_ = (Ψ_LR_, Ψ_FH_, Ψ_DIAG_) = (0.5,0.5,0)] to the idealized pace [Ψ_B_ = (Ψ_LR_, Ψ_FH_, Ψ_DIAG_) = (0.5,0,0.5)] for the contusion (C). The thin black lines show the extension of the lines up to a distance of 1 from their reference points Ψ_&_; this is also the maximal distance from the refence point for which the projections were calculated. l, left; r, right; f, forelimb; h, hindlimb.

As we described above, rats that recovered locomotion after lateral hemisection injuries lost high-speed left–right synchronous gaits, but could still use trot, canter, and gallops that were similar to those of intact rats, although with a contralateral lead-limb bias during asymmetric gaits. The distributions of phase differences of hemisected rats (Figure 8B; Figure 9B) corroborate this finding as they appear to be a subset of those of the intact rats. Indeed, 73.77% of the variance of phase differences could be explained by the same line between trot and half-bound gallop as in intact rats (R^2^ per rat: median=0.677, inter quartile range: 0.608–0.702). This also provides an explanation of why the homolateral phase differences ipsilateral (right) and contralateral to the lesion appear to be affected differently by the lesion. Across all speeds and gaits, the ipsilesional homolateral phase differences exhibited significantly lower values compared to before the injury (*χ*^2^_1_ =5.980, p=0.015; Supplementary Material), while the contralesional ones remained unchanged (*χ*^2^_1_ =0.000, p=0.989). Yet, when comparing these phase differences only for trot and gallop with a left lead limb, no significant changes were detected for either homolateral phase differences (gallop, ipsilesional: *χ*^2^_1_ = 0.004, p=0.951, contralesional: *χ*^2^_1_ = 0.704, p=0.401; trot, ipsilesional: *χ*^2^_1_ = 0.631, p=0.427; contralesional: *χ*^2^_1_ = 0.003, p=0.954). Thus, both homolateral phase differences after hemisection injury remained appropriate for the specific gait.

At the same time, the line between trot and half-bound gallop does not fit well to the distribution of phase differences after contusion injury (R^2^ across animals: 0.446, R^2^ per rat: median=0.419, inter quartile range: 0.221–0.452; Figure 9C). The contused animals did not exhibit speed-dependent changes of the phase differences (Figure 8C). Phase differences close to trot (left– right and homolateral alternation, diagonal synchronization) were most common and thus were concentrated around alternation for both forelimb and hindlimb pairs, along with substantial variability of the homolateral and diagonal phase differences spanning the whole cycle (Figure 8C; Figure 9C).

The interlimb coordination of a quadruped is fully defined by three phase differences and our data show that, after a contusion injury, the normalized left–right phase differences of the forelimbs (ф_FLR_) and the hindlimbs (ф_HLR_) have been controlled to be close to alternation:

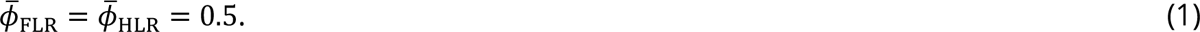

Thus, only one degree of freedom remains. Since the forelimb normalized left–right phase difference is related to the homolateral (ф_HL_) and diagonal phase differences (ф_DIAG_) by

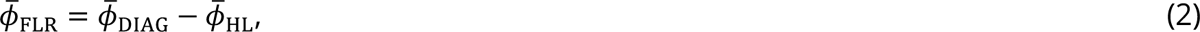

this remaining degree of freedom follows (obtained by substituting equation 1 into equation 2):

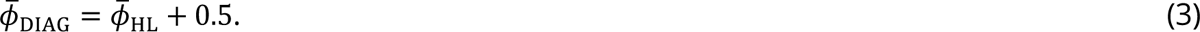

Both trot and pace (Table 1) satisfy equations 1–3 and the remaining degree of freedom can be interpreted as the line connecting these gaits. Indeed, the distribution of phase differences was oriented along the line between trot and pace [Figure 9C; R^2^ across animals: 0.840, R^2^ per rat: median=.819, inter quartile range: 0.754–0.837; AICw=1.000 (compared to the line between trot and half-bound gallop)]. Since lateral-sequence and diagonal-sequence gaits (walk or amble, depending on the duty factor) also lie along this line (Table 1), it includes all gaits expressed after contusion. This distribution of phase differences could be the result of strong left–right coupling of the forelimbs and hindlimbs that maintained left–right alternation together with relatively weak coupling between the forelimbs and hindlimbs. To investigate this further, we analyzed step-to-step variability in the following.

### Contusion and hemisection injuries differentially affect interlimb coupling

The central neuronal control of interlimb coordination is presumed to be mediated by commissural and long propriospinal neurons (Talpalar et al., 2013; Kiehn, 2016; Ruder et al., 2016; Danner et al., 2017; Flynn et al., 2017; Frigon, 2017). These neurons couple the four rhythm generators, each controlling one limb. Changes in the coupling strengths are expected to be reflected in the variability of the phase differences between the corresponding pairs of rhythm generators and consequently between the corresponding limb pairs. Thus, by analysis of the variability of phase differences, we can gain insight into the altered connectivity between the rhythm generators. To quantify the variability of the phase differences between each pair of limbs, we calculated their mean deviation from the circular exponential moving average for each bout (see Methods for details).

The mean deviation from the moving average was chosen as it accounts for changes in phase differences due to gait transitions. This can be seen in two exemplary bouts in Figure 10A: in the bout of the contused animal (Figure 10A1), the homolateral and diagonal moving averages follow their phase differences, but the deviation persists throughout the bout as the phase differences keep changing and the gait never stabilizes. Yet, in the bout of the intact animal (Figure 10A2), the left–right and diagonal moving averages only deviate from their phase differences during the gait transitions and converge at the end of the bout. Thus, while still an imperfect measure, ordered gait transitions result in lower mean deviations from the moving average than persistent or stochastic changes.

**Figure 10.**
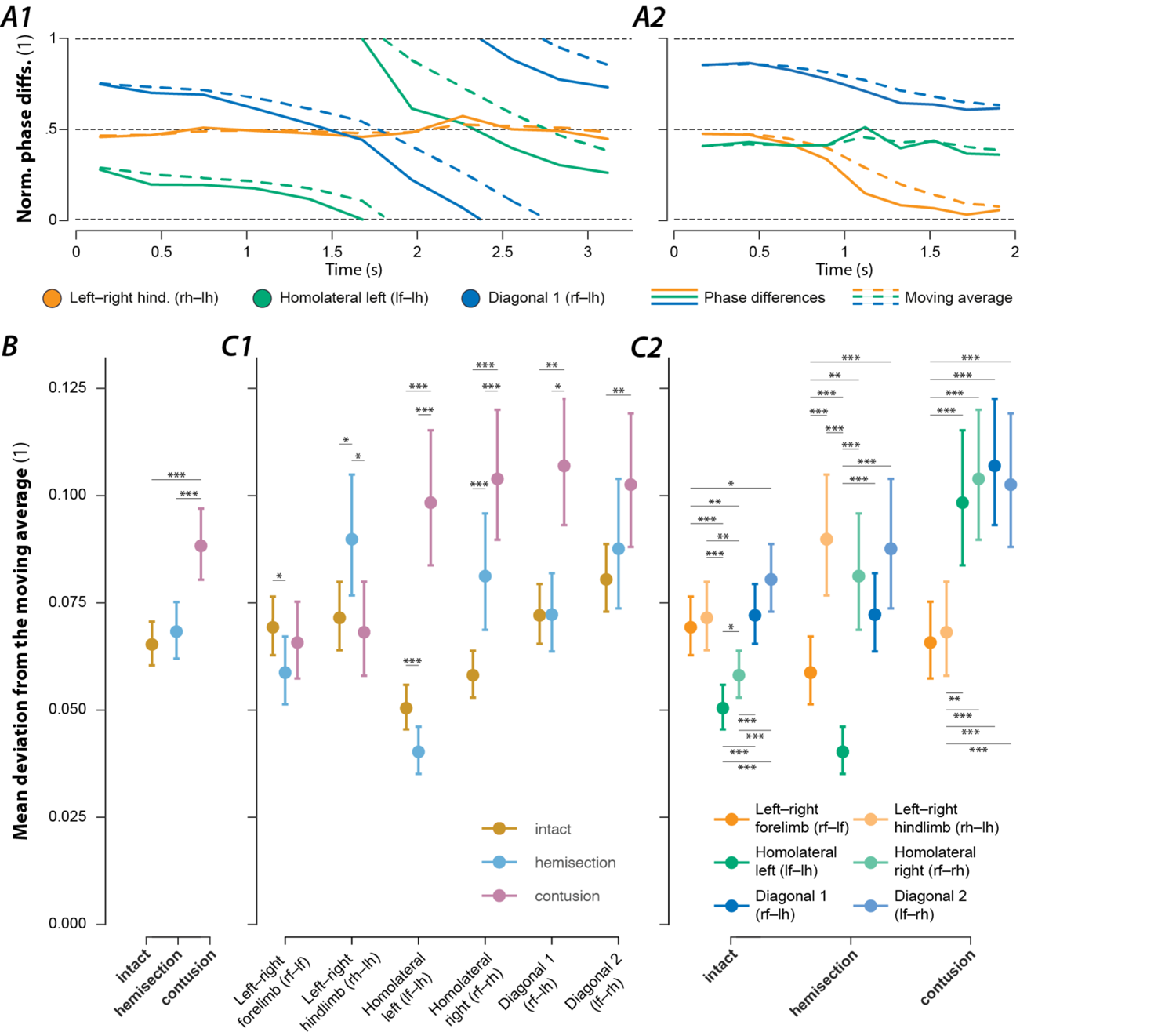
Variability of interlimb coordination in intact rats and after hemisection and contusion injury. (A) Two examples (A1: contusion, A2: intact) of normalized phase differences (solid lines) and their moving averages (dashed lines). (B) Marginal means of the mean deviation from the moving average of the phase differences of intact animals and following hemisection or contusion injury. (C) Means of the deviations from the moving average of each phase differences for intact rats and rats after recovery from hemisection and contusion injuries. A generalized linear mixed model with injury type (intact, hemisection, contusion) and phase difference (left–right hindlimb, left–right forelimb, homolateral and diagonal on each side) as well as their interaction effect as fixed effects, a per-animal random offset, and a full factorial dispersion model was calculated for the deviation of the moving average. A logit link function and beta distribution were used to account for the non-normality of the residuals. All fixed and interaction effects were significant (injury type: F2,1641=39.741, p<0.0001; phase difference: F5,1641=23.863, p<0.0001; interaction effect: F10,1641=13.730, p<0.0001). Asterisks denote significant post-hoc tests between injury types (C1) and phase differences (C2). l, left; r, right; f, forelimb; h, hindlimb. *, p < 0.05; **, p < 0.01; ***, p < 0.001. Error bars denote 95% confidence intervals.

The mean deviation from the moving average of the phase differences across all pairs of limbs (Figure 10B) was significantly higher after contusion injury compared to pre-injury or hemisection injury. Yet, hemisection injury did not result in significantly different values compared to pre-injury.

In intact rats, the mean deviations from the moving average of both homolateral phase differences were significantly lower than those of the left–right and diagonal phase differences (Figure 10C2). This suggests that coupling of the homolateral neuronal circuits on each side of the spinal cord was stronger than the coupling of left and right circuits within or between enlargements.

Figure 10C shows that the mean deviation from the moving average of the forelimb and hindlimb left–right phase differences did not significantly change after contusion injury. Yet, the mean variability of all phase differences involving a forelimb and a hindlimb significantly increased. Furthermore, the mean variability of each left–right phase difference was significantly lower than that of all homolateral and diagonal phase differences (Figure 10C2). This suggests that after contusion injury, coupling between the left and right limbs within each enlargement remained at (or recovered back to) the pre-injury levels while the coupling between the lumbar and cervical circuits was significantly reduced. An alternative suggestion is that inter-enlargement circuitry introduces variability to the left–right phase differences at each girdle, that is reduced following a partial loss of long propriospinal axons.

The changes in the deviation of the moving average of the phase differences were more complex following a hemisection injury (Figure 10C). The mean variability of the forelimb left–right phase differences significantly decreased after injury, while that of the hindlimb left–right phase differences significantly increased. As for homolateral phase differences: the mean variability significantly increased on the side ipsilateral to the injury (right side) and significantly decreased on the side contralateral to the injury. The phase differences for the diagonal limb pairs did not significantly differ compared to before the injury. These results suggest that the neuronal connections that directly coupled the lumbar and cervical circuits prior to injury were compensated for by some spared or potentially new neuronal connections after hemisection.

## Discussion

### Speed-dependent gait expression in intact rats

For this study we utilized animals trained in a 3-m long tank using food incentives to rapidly locomote back and forth on a high-friction surface (sylgard), which provided an ideal setting to analyze high-speed gaits and speed-dependent gait transitions. Rats started from rest, quickly started locomoting, and transitioned from the alternating gaits of walk and trot through the transitional gait of canter to the high-speed non-alternating gaits of transverse gallop and half-bound gallop, finally arriving at the synchronous gait of bound. Using this strategy, we confirmed and extended the findings of several previous studies that characterized speed-dependent gait transitions using other experimental settings with more restricted animal locomotion or a narrower range of speeds and gaits. For example, Cohen and Gans (1975) reported walk, trot, canter, and gallop across wide range of speeds, but did not observe fast enough speeds for the animals to express bound. They reported a top speed of 125.9 cm/s and Heglund and Taylor (1988) of 123 cm/s, whereas uninjured animals in the present study achieved an average maximal speed of 188±8.85 cm/s.

Confirming and extending these and other previous studies, we observed that a lateral-sequence walk was expressed at the slowest speeds. The trot expressed was actually a running trot with a duty factor of less than 0.5 and a brief suspension phase between the ground contacts of the two diagonal limb pairs (Hildebrand, 1976, 1989). Others have reported both walking and running trots, where the walking trot appeared only at slow speeds (Górska et al., 1999), thus it could have been absent in our experiments due to our training strategy and apparatus. During gallop (and even half-bound gallop), our rats rarely exhibited a suspension phase. While this confirms previous studies in rats (Cohen and Gans, 1975), other species (Hildebrand, 1977, 1989), including mice (Bellardita and Kiehn, 2015; Lemieux et al., 2016), often exhibit a significant suspension phase during gallop. However, during bound (the fastest gait) two suspension phases occurred—a longer one and a shorter one, after the lift-off of the hindlimbs and of the forelimbs, respectively.

Finally, we observed a three-beat canter that resembled the same gait in horses (Harris, 1993; Bertram, 2016) and occurred as a transitional gait between trot and gallop, where one pair of diagonal limbs moved in-phase as during trot, while the other was offset by a quarter to a third of a cycle as during gallop. Cohen and Gans (1975) described canter to occur while speeding up during trotting before transitioning to a full gallop, but it is not clear if they observed the same three-beat gait, since their definition of canter only relied on left–right asymmetry. Overall, these characteristics likely reflect key control mechanisms and their interaction with the plexiglass tank and the high-friction sylgard walking surface, and the training strategy used to incentivize rapid transitions to high-speed gaits.

### Characteristics of locomotion following recovery from thoracic lateral hemisection and contusion

Following recovery from spinal cord injury, we observed several changes of gait parameters that are in accordance with previous literature. For example, thoracic contusion injury results in a decrease in locomotor speed (Hamers et al., 2001) and an increase in duty factor (due to the decrease of swing duration, Beare et al., 2009; Redondo-Castro et al., 2013). Other changes include a decreased use of walk with a lateral sequence compared to other walk patterns (Hamers et al., 2001) and an increased variability of homolateral and diagonal phase differences (Kloos et al., 2005). Our results also showed the emergence of the gaits pace and amble, the latter of which occurred at higher walking speeds and frequencies, and might have been missed with protocols that investigate a restricted range of speeds. Following a lateral hemisection injury, rats develop compensation mechanisms to help stabilize locomotion (Webb and Muir, 2002) such as left–right asymmetry in duty factors (Webb and Muir, 2002; Ham et al., 2019) and an increased base of support (Kunkel-Bagden et al., 1992). More dispersed diagonal phase differences have also been reported (Ham et al., 2019), which our analysis did not show. This discrepancy was likely due to the animals attempting to transition to gallop and an artifact of the analysis of the dispersion of phase differences relative to the grand mean rather than to the moving average as performed here (cf. Figure 10).

Importantly, our analysis showed that the modulation of stride length and step cycle duration with locomotor speed did not differ between pre-injury and post-recovery locomotion, neither for contusion nor hemisection injuries. This likely reflects the robust retention of control mechanisms within modified spinal circuitry in the context of the experimental training and apparatus employed, which provided essentially optimal conditions for the expression of stable high-speed gaits.

Furthermore, it is also possible that hypertonia, hyperreflexia, and clonus influenced intra-limb coordination and consequently locomotor function. While these signs are common observations in humans after spinal cord injury, they are rare in rodents with all but the most severe contusions or anatomically complete transections. Following moderate T2 or T10 contusions, intralimb coordination (inter-joint coordination) is only temporarily disrupted (Shepard et al., 2021), and likely did not have a major impact on the observed locomotor behavior.

### What causes the reduction of locomotor speed following hemisection and contusion?

Spinal cord injury disrupts communication across the injury site and, hence, affects ascending and descending pathways to and from supraspinal centers as well as ascending and descending propriospinal pathways. In addition, the injury affects primary and secondary somatosensory pathways. These pathways are involved in various functions and their disruption caused by contusion and hemisection injuries result in different anatomical changes, thus our results can be used to infer functional-anatomical relationships resulting from partial or full loss of specific pathways.

After a mid-line spinal contusion injury (mild-moderate severity; as applied here), the dorsal columns (somatosensory axons) and the corticospinal tracts are significantly disrupted. In addition, there is a mild disruption of lateral and ventral white matter tracts (Kim et al., 2012). Despite these significant anatomical disruptions, after recovery these animals were able to demonstrate stable locomotion over a wide range of speeds with essentially normal speed-dependent changes in stance and swing durations, duty factor, and right-left phase differences of both limb pairs. However, the injury disrupted forelimb–hindlimb coordination and induced disorganized gait patterns, leading to an inability to increase speed and to express the high-speed gaits of gallop and bound. This disruption of forelimb–hindlimb coordination likely reflects weakened coupling of the limb-specific neuronal circuits interconnected via long-propriospinal neurons and/or by a reduction of descending drive to the lumbar spinal cord.

In contrast to the contusion injury, the hemisection injuries selectively remove all the ascending and descending axons on one side including supraspinal, propriospinal, and sensory pathways. Importantly, it leaves one side completely intact, and thus the strong bilateral distribution of pathways via commissural interneurons within the intact enlargements allowed for remarkably good functional recovery. While forelimb–hindlimb coordination was affected by the hemisection, the homolateral and diagonal phase differences were generally within normal ranges. Yet, the hemisection induced a lead-limb bias, increased variability of left–right hindlimb coordination (phase differences) and caused asymmetric coupling of forelimbs and hindlimbs. We speculate that these asymmetries prevent the animal from achieving the synchronization of the left and right hindlimbs necessary for the high-speed gaits of half-bound gallop and bound. It is also possible that other biomechanical or neuronal factors, such as the loss of local trunk control ipsilateral to the injury or the loss of cortico- and reticulospinal inputs to the lumbar spinal cord ipsilateral to the injury, caused a reduction of the propulsive force that the animals could generate. Computer simulations are necessary to further delineate potential mechanisms.

### Distribution of phase differences and their variability give insights into neuronal coupling and plasticity

In intact rats, phase differences of expressed steps were distributed as a continuum along a line in phase space (Figure 9A). This distribution included trot, gallop, half-bound gallop, and bound without clear borders separating the distinct gaits or obvious clustering of datapoints. There were numerous steps in-between the areas representing strongly defined gaits. One interpretation of these results could be that discrete gaits (as defined by interlimb coordination) do not exist but resulted from the analysis or experimental conditions used previously (i.e., the setup or apparatus biases the animals to a few gaits and speeds). Extending that concept, the continuous distribution of phase differences that we observed could result from our specific experimental setup, which biased the animals towards constantly changing speed and gait, suggesting that gait changes occurred gradually and involved transitional patterns, as also observed in mice in response to perturbations (Vahedipour et al., 2018). Interestingly, we previously observed a distribution of left–right phase differences across the entire possible range for the hindlimbs following synaptic silencing of the L2-to-L5 propriospinal interneurons (Pocratsky et al., 2017) and for both the forelimbs and hindlimbs following synaptic silencing of the L2-to-C6 long-ascending propriospinal neurons (Pocratsky et al., 2020). Thus, in either case, spinal cord injury or synaptic silencing, the distribution of phase differences allows us to speculate about the neuronal control of interlimb coordination.

The orientation along a line suggests that the neuronal circuitry constrains the direction of possible changes of interlimb coordination and that a single mechanism underlies the transition between the gaits along this line. This is in accordance with our computational models of interlimb coordination and speed dependent gait expression in mice (Danner et al., 2016, 2017; Ausborn et al., 2021; Zhang et al., 2022), where the transition from the alternating gait trot to the (quasi-)synchronous gaits gallop and bound was caused by a shift in the balance in the activity of commissural and long propriospinal interneurons that promoted left–right alternation or diagonal synchronization to those that promoted left–right synchronization. It is important to note that the transition from trot to gallop in the computational models exhibited hysteresis, which is common in quadrupeds (Heglund and Taylor, 1988). However, it is not clear whether such a multistable transition could underly the current observations; a detailed analysis of model dynamics during speed changes and gait transitions would be necessary.

Interestingly, while rats that recovered locomotion after a lateral hemisection lost the high-speed synchronous gaits and always used the same lead-limb for asymmetric gaits, the distribution of their phase differences (Figure 9B) still fell on the same line as pre-injury. This suggests that hemisected animals employed the same mechanism for gait control as they did before the injury, but the range of gaits was restricted by the loss of one-half of the ascending and descending, homolateral and commissural propriospinal neurons. On the other hand, the distribution of phase differences exhibited by rats that recovered locomotion after the contusion injury (Figure 9C) revealed that the expressed behavior, including the new gaits that were not expressed pre-injury, could be explained by maintenance (and perhaps strengthening) of left–right alternation of forelimbs and hindlimbs together with weaker forelimb–hindlimb coupling (see also Figure 10B). Thus, the underlying mechanisms of locomotor control following recovery from hemisection and contusion injury likely differ.

### Recovered locomotion

As discussed above, the analysis of the variability of phase differences between all limb pairs (Figure 10) provides insight into the organization of spinal circuitry following recovery. Low variability of phase difference represents strong coupling between the specific pair of limbs, and high variability represents weak coupling. We observed interlimb coupling strengths after lateral hemisection injury (Figure 10B) that suggest that the sprouting and reorganization known to occur after hemisections (Takeoka et al., 2014; Filli and Schwab, 2015) plays a role in the recovery of locomotion. The increase in homolateral coupling contralateral to the injury as compared to pre-injury suggests strengthening of homolateral long-propriospinal connections in the intact hemicord. Also note that, while homolateral coupling ipsilateral to the injury was weakened compared to pre-injury, it was still stronger than following an incomplete contusion injury, which only damaged the same neuronal pathways partially. Together with the observation that diagonal coupling was not different to pre-injury, this suggests that new propriospinal connections were formed that could compensate for the anatomical hemisection. Such connections could include long-propriospinal neurons that cross the midline twice as well as more complicated oligo-synaptic pathways (Takeoka et al., 2014; Filli and Schwab, 2015).

Furthermore, the presence of 1-to-1 coupling between all limbs implies that the brainstem drive to the lumbar rhythm generator ipsilateral to the hemisection injury recovered. This could be due to reorganization of descending reticulospinal pathways (Ballermann and Fouad, 2006; Filli et al., 2014; Zörner et al., 2014; Filli and Schwab, 2015)—e.g., through sprouting of contralateral descending fibers crossing over the midline or forming synapses to contralateral lumbar commissural interneurons—and compensation by primary afferent feedback signals and intraspinal reorganization (Barriere et al., 2008; Cohen-Adad et al., 2014; Takeoka et al., 2014; Takeoka and Arber, 2019). Similar observations following recovery from the mild-to-moderate contusion injury could be explained by a combination of partial sparing of reticulospinal neurons, reported to be common with such injuries (Filli et al., 2014; Fouad et al., 2021), together with their reorganization and sprouting, and compensation by afferent feedback.

Recently (Shepard et al., 2021), we showed that silencing L2-to-C6 long-ascending propriospinal neurons after recovery from mid-line contusion injuries, similar to those employed in the current study, actually resulted in an improvement of functional locomotion. Silencing induced a modest increase in plantar paw placement (as opposed to stepping on the curled toes or dorsum of the paw) and a dramatic improvement in left–right alternation of the hindlimbs (variability of phase differences was significant reduced). We suggest that the partially spared long-ascending propriospinal neurons could allow for a range of left–right hindlimb phase differences after recovery from injury and that removal of that ascending component leads to an improved or strengthened left– right coupling. The variability in phase-differences returned when synaptic silencing ceased. Thus, even in the absence of anatomical sprouting, altering the relative coupling strength between and within enlargements can impact gait characteristics.

It is worth a reminder that our experimental design focused on uninjured and recovered locomotion, not on the changes occurring during the recovery process. Thus, our observations cannot be interpreted to reflect the pre-injury circuitry minus that directly damaged, but instead it reflects the capacity/operation of the spinal cord following the full amount of spontaneous recovery. To draw concrete conclusions about reorganization of spinal pathways, additional anatomical and computational studies are necessary.

### Analysis of overground locomotion with a wide range of speeds uncovers specific deficits resulting from spinal cord injury

Hemisected rats were able to locomote well at slow to intermediary speeds. Differences to pre-injury in slow speed, alternating-gait locomotion were relatively minor: the duty factor was increased and exhibited a slight left–right asymmetry for the hindlimbs, and a slight shift of left–right phase differences occurred during trot. Yet, at intermediary speeds, hemisected rats only used gallops with a left lead (contralateral to the injury); the high speed (quasi-) synchronous gaits (half-bound gallop and bound) were not expressed, and both the maximum and average speeds were reduced. Thus, assessment strategies that limited analysis to slower speeds would have missed several important changes in the locomotor behavior after recovery from injury.

## Methods

### Ethical approval

All experiments were performed in accordance with the Public Health Service Policy on Humane Care and Use of Laboratory Animals, and with the approval of the Institutional Animal Care and Use Committee (IACUC, protocol number 19644) at the University of Louisville.

### Animals, spinal cord injury and experimental setup

A group of 17 Adult female Sprague-Dawley rats (225–250g) were housed in pairs (with one cage of three) under a 12-hour light/dark cycle with food and water provided *ad libitum*. All 17 animals were assessed pre-injury and then randomly divided into two groups that received either hemisection injuries (nine rats) or contusion injuries (eight rats). Three animals with hemisection injuries were removed from the study due to autophagia of toes on their hind paw, leaving a final group size of six. One animal with a contusion injury was removed because the contusion was not delivered appropriately, and it was walking immediately upon recovering from the anesthesia leaving a final group size of seven. For each animal and injury condition, 5–15 locomotor bouts, with 9.1±1.92 full step-cycles each, were analyzed.

For contusion and hemisection injuries animals were anesthetized (ketamine:xylazine, 80 mg/kg:4 mg/kg; Henry Schein Animal Health; Akorn Animal Health), the lower thoracic spinal column was surgically exposed using a dorsal approach and a T9 laminectomy was performed. The spine was immobilized at the T8 and T10 levels using custom-built stabilizers and the injuries were delivered as previously described for a mild-moderate (12.5 g-cm) mid-line contusion injury at T10 using an NYU Impactor device (Magnuson et al., 2009) or for a lateral hemisection using micro-scissors (Beaumont et al., 2006). Post-injury care included antibiotics, analgesics, daily manual bladder expression and supplemental fluids as previously described (Magnuson et al., 2009).

The lateral hemisection used results in complete loss of ascending and descending axons, including the dorsal column and corticospinal tract, on one side of the spinal cord. In contrast, the mild-moderate contusive injury results in essentially complete loss of the dorsal columns and corticospinal tract bilaterally and approximate 50% loss of dorsolateral, lateral, and ventrolateral white matter bilaterally. In most animals, the ventral columns were slightly more damaged than the lateral columns. Images/imaging of spinal cords from animals with similar injuries can be found in numerous publications (Kim et al., 2012; Beaumont et al., 2006; Basso et al., 1995).

Locomotion was assessed in a 3-m long Plexiglas tank where the walking surface was coated with sylgard to provide good grip. The rats were trained, two or three at a time, using food treats, to traverse the length of the tank, and this training strategy resulted in all animals utilizing left–right synchronization of the hindlimbs on at least some of the recorded passes. Recordings were made using four high-speed (200 Hz) video cameras placed to capture the ventral view of the animal (Figure 1A). Custom software was used to stitch the videos together. Foot placement timing was recorded and analyzed as described below. Following multiple baseline assessments animals were divided into two groups, as described above, and given either lateral hemisection or midline contusion injuries. They were allowed to recover for at least 4 weeks at which time their locomotor function reached a plateau based on the standard BBB Open Field Locomotor Scale with mean scores of 15.7±1.9 (contused) and 18.3±0.8 (hemisected) on the scale out of 0–21 at day 42 post-injury (Basso et al., 1995). The animals were then re-introduced to the long-tank set-up and trained again (at least 2 at a time) to traverse the tank for a food treat. Multiple complete passes were then recorded for each animal. We did not assess, train, or otherwise manipulate the animals at early post-injury time points, beyond normal post-operative care, to avoid any potential functional impact.

### Data analysis

The locomotor period or step-cycle duration was defined as the duration between the midstance time (halfway point between onset and offset of stance) of two consecutive steps of the reference limb (left hindlimb, if not noted otherwise); the frequency was defined as the reciprocal of the period. For each limb, stance and swing durations were calculated as the duration between onset and offset of stance and swing, respectively; the duty factors were calculated as the stance duration divided by the period; the stride lengths were calculated as the Euclidean distance between two consecutive ground contact points; and the instantaneous speeds were calculated as the average of the stride lengths of each limb divided by the period. Maximal speed was operationally defined as the 95^th^ percentile of the instantaneous speeds. Limb-specific parameters were then averaged for all four limbs or for the forelimbs and hindlimbs separately. Normalized phase differences for each pair of limbs were calculated as the delay between the midstance times divided by the period. Phase differences (in radians) were calculated by multiplying the normalized phase differences by 2π. Normalized phase differences are denoted by a bar over the variable.

#### Gait classification

For each step cycle, the gait was determined based on phase differences and the duty cycle. The trailing limb of a pair was defined as the limb that strikes the ground first; the one that strikes second was defined as the lead limb. First, each step was compared to all idealized one, two, three, and four beat gaits with equal phase-shifts between the beats (Table 1). A step was assigned to belong to the gait whose idealized representation was closest in phase-space. Then, pairs of gaits that only differ based on their lead limb were grouped together and walking and running gaits were separated based on whether the duty factor was larger (walking gaits) or smaller than 0.5 (running gaits). Table 1 shows the normalized phase differences of all idealized gaits and the subsequent classification based on duty factor. Note that an additional two-beat gait with homolateral and diagonal normalized phase differences of 2/3 was introduced to better account for the fore–hind asymmetry common in bound (Hildebrand, 1977; Bellardita and Kiehn, 2015).

To calculate the distance between each step and the idealized gaits, both steps and gaits were represented as points (Φ and Ψ) in the three-dimensional space consisting of hind left–right, homolateral, and diagonal phase differences (the trivial phase difference between the left hind limb and itself was omitted), and the distance was calculated as:

#### Gait transitions

A gait transition was defined as a step of one gait being followed by a step of another gait. Occurrences for each ordered pair of two gaits were calculated and assembled into a matrix A, where *a_i,j_* is the number of steps of gait *i* that were followed by gait *j*. Conditional probabilities of gait transitions were obtained by normalizing each element by the sum of elements in that row (shown in Figure 2H) and frequencies of gait transitions were obtained by normalizing each element of A by the total number of gait transitions (shown in Figure 2I).

#### Variability of phase differences

The variability of the phase differences was quantified by calculating the deviation of the circular exponential moving average. The moving average of the phase differences reflects a weighted average of the most recent steps rather than of all steps and was iteratively calculated by *S*_0_ = ф_0_, and

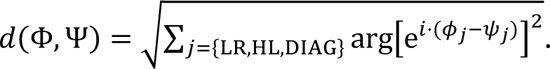

where *n* is the index for the step cycle within a given bout, ф is the sequence of phase difference in radians, and α is the smoothing factor. α and the cutoff frequency *f*_c_ are related as.

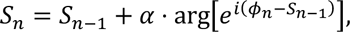

For each bout, the mean deviation of a phase difference from its moving average was calculated as:

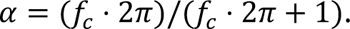

Note the normalization factor π. The cutoff frequency *f*_c_ was set to 0.125 step cycles.

#### Statistics

Statistical analysis was performed using lme4 1.1.31 (Bates et al., 2015) and glmmTMB 1.1.5 (Kristensen et al., 2016; Brooks et al., 2017) for R 4.2.2 (R Core Team, 2020) and using SciPy 1.9.1 (Virtanen et al., 2020), lmfit 1.0.3 (github.com/lmfit/lmfit-py), and NumPy 1.23.2 (Harris et al., 2020) for Python 3.10 (Python Software Foundation, Wilmington, Delaware, USA). The specific statistical models and methods used are listed in the following. For linear and generalized-linear mixed effect models, model assumptions were confirmed both by visually inspecting the quantile-quantile plots and histograms, and by performing distribution, dispersion, outlier, and quantile deviation tests using the DHARMa R package (Hartig, 2022). If model assumptions were not met, link functions and error distributions that conform with the assumptions were selected. Linear mixed models were fit using lemr of the lme4 R package; generalized linear mixed models, or linear mixed models that require a dispersion model, were fit using glmmTMB. Mixed-model designs were chosen to adequately account for repeated measures and within as well as between animal variances. Marginal means were estimated using emmeans and reported as mean ± standard error. To account for multiple comparisons, Tukey’s method was applied to all post-hoc tests.

For the comparison of locomotor speed, step frequency, stride length, and duty factor between the different gaits of intact rats (Figure 1C–F) as well as of hemisected (Figure 5B–E) and contused rats (Figure 7B–E), separate linear mixed models with gait as a fixed effect and a by-subject random intercept were calculated. Degrees of freedom were adjusted using the Kenward-Roger method to account for violations of sphericity.

For the comparison of locomotor speed, step frequency, stride length, duty factor, and maximal speed (Figure 2B–F) between pre-injury (intact) rats and rats following either hemisection or contusion (injury type), separate mixed models with injury type as the fixed effect, a per-rat random intercept and a random slope of injury type, as well as a full-factorial dispersion model were calculated using glmmTMB. For speed, frequency, and maximal speed, linear mixed models were calculated; for stride length, a log link was used; for duty factor, a Gamma error distribution and a logit link function was used.

To assess differences of left–right forelimb and hindlimb phase differences during trot between intact and hemisected animals (Figure 6), separate linear mixed models with injury type as the fixed effect, a per-rat random intercept and slope of injury type, as well as a full-factorial dispersion model were calculated using glmmTMB.

For comparison of the mean deviations of phase differences from their moving average between injury type and limb pairs (Figure 10), a generalized linear mixed model with injury type (intact, hemisection, contusion) and phase difference (left–right hindlimb, left–right forelimb, homolateral and diagonal on each side) as well as their interaction effect as fixed effects, a per-animal random offset, and a full factorial dispersion model was calculated. A logit link function and beta distribution were used to account for the non-normality of the residuals.

To assess speed-dependent modulation of gait characteristic of intact and injured rats (Figure 3), linear and non-linear regressions were performed on a per-animal and injury type basis using lmfit. Group averages were calculated using the average of the individual regression results and the confidence interval (CI) was calculated using the delta method:

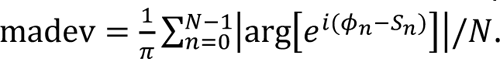

where f is the regression function and ∇f it’s gradient, µ_β_ the average parameter estimates, ν the instantaneous locomotor speed, and *Cov*_β_ the average variance-covariance matrix of the individual parameter estimates. Linear regressions (*f*(β, ν) = β_0_ + β_1_ * ν) were calculated for the stride length, forelimb and hindlimb duty factor and forelimb and hindlimb swing duration; non-linear regressions with an inverse function (*f*(β, ν) = β_0_ + β_1_ * ν^-1^) were calculated for the step cycle duration and forelimb and hindlimb stance duration.

To quantify the distribution of individual steps on a continuum in phase-space (Figure 9), we calculated the projection λ_i_ of the *i*-th step Φ_+_ from a vector defined by two average or idealized gaits (Ψ_A_ and Ψ_B_):

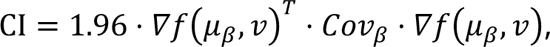

Φ_+_, Ψ_B_, and Ψ_C_ are points in the three-dimensional space of left–right hindlimb, homolateral, and diagonal phase-differences. To account for the circular nature of phase differences, the projections were recalculated by shifting the reference position by ±2πu^:

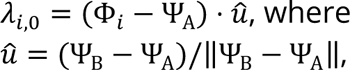

and selecting the projection (λ_+,7_, λ_+,!_, λ_+,/!_) for which the distance to the line is minimal

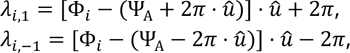

Furthermore, values of λ_+_ were restricted to |λ_+_| ≤ 2π:

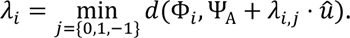

To assess the model fit, the coefficient of determination was calculated per animal and across animals:

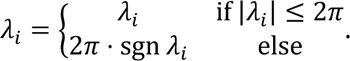

where µ_F_ is the grand circular mean of Φ. To compare two alternative models, additionally the Akaike Information Criterion (AIC) was calculated for each model:

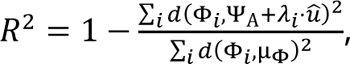

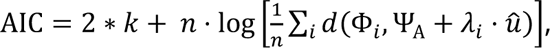

where *k* is the number of parameters and *n* the sample size. AIC weights, representing the relative likelihood of a model compared to the alternative models (Burnham and Anderson, 2002), are reported. A line between the average trot 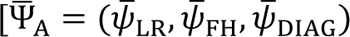 = (0.49,0.47,0.98)] and the average half-bound gallop of intact animals 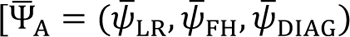 = (0.94,0.60,0.36)] was used; for the contusion, the fit of this line was compared to that of a line between the idealized trot 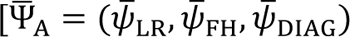 = (0.5,0.5,0)] and pace 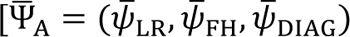 = (0.5,0.5,0)] (Table 1).

## Additional information

### Data availability statement

All raw data and source code to analyze the data, perform statical tests, and create figures are available at https://github.com/dannerlab/rat-sci-locomotion.

### Author contributions

Conceptualization, S.M.D, D.S.K.M.; Resources, D.S.K.M.; Data curation, S.M.D.; Formal analysis, S.M.D.; Funding acquisition, S.M.D., D.S.K.M.; Validation, S.M.D., C.T.S, C.H., N.A.S., I.A.R., D.S.K.M.; Investigation, C.T.S, C.H.; Visualization, S.M.D., N.A.S.; Methodology, S.M.D., C.T.S, C.H., N.A.S., I.A.R., D.S.K.M.; Writing—original draft, S.M.D., D.S.K.M.; Writing—review & editing, S.M.D., C.T.S, C.H., N.A.S., I.A.R., D.S.K.M.

